# A neural progenitor mitotic wave is required for asynchronous axon outgrowth and morphology

**DOI:** 10.1101/2021.03.10.434802

**Authors:** Jérôme Lacoste, Hédi Soula, Angélique Burg, Agnès Audibert, Pénélope Darnat, Michel Gho, Sophie Louvet-Vallée

**Author notes:** co-last authors (Michel Gho, Sophie Louvet-Vallée).

## Abstract

Spatiotemporal mechanisms generating neural diversity are fundamental for understanding neural processes. Here, we investigated how neural diversity arises from neurons coming from identical progenitors. In the dorsal thorax of *Drosophila*, rows of mechanosensory organs originate from the division of sensory organ progenitor (SOPs). We show that in each row of the notum, a central SOP divides first, then neighboring SOPs divide, and so on. This centrifugal wave of mitoses depends on cell-cell inhibitory interactions mediated by SOP cytoplasmic protrusions and Scabrous, a secreted protein interacting with the Delta/Notch complex. Furthermore, when the mitotic wave was abolished, axonal growth was more synchronous, axonal terminals had a complex branching pattern and fly behavior was impaired. We show that the temporal order of progenitor divisions influences the birth order of sensory neurons, axon branching and impact on grooming behavior. These data support the idea that developmental timing controls axon wiring neural diversity.

## INTRODUCTION

It is commonly accepted that nervous system development relies on the precise spatio-temporal regulation of gene expression in neural progenitors ^1^. However, little is known about how neural diversity can be generated among neural progenitors that are homogenously specified ^2^, for example, in neurons that connect peripheral sensory organs with the central nervous system ^3^.

Microchætes are peripheral mechanosensory organs on the thorax of *Drosophila melanogaster*. These organs arise from sensory organ progenitor cells (SOPs) which are selected among G2 arrested cells of proneural clusters during pupal stage ^4,5^. In the dorsal region of the thorax (notum), SOPs appear sequentially from five parallel proneural rows: first rows 5, followed by rows 1 and 3 and finally rows 2 and 4 ^5,6^. SOP selection involves Notch-dependent lateral inhibition from 6 to 12 h after pupal formation (APF). Then, at 16.5 h APF, SOPs resume the cell cycle and undergo a sequence of divisions to generate a neuron and three other non-neural support cells (the shaft, the sheath and the socket cells) to form each organ ^7^. During organogenesis, the bipolar neurons project their dendrite toward the base of the bristle, and their axons toward the ventral thoracic ganglion. The axons enter the ganglion through the posterior dorsal mesothoracic nerve root and extend branches anteriorly and posteriorly ^8^. The resulting arborization of the microchæte axon projections is variable. Their complexity is not correlated to the position of the organ but to the time at which rows appear. Thus, the earlier the microchaetes developed, wherever they were positioned, the more branched they were, arborizing over a greater area within the neuropil ^8,9^. In adult flies, mechanical stimulation of thoracic bristles induces a cleaning reflex in which the first or third leg sweeps the stimulated area. This reflex requires the correct connectivity of the bristle neurons ^8,10^.

SOP distribution in the notum depends on N-signaling pathway as well as modulators, such as Scabrous (Sca). Scabrous is a secreted fibrinogen like protein that interacts physically with N and Dl and modulates their activities ^11^, The precise mechanism by which Sca regulates the N-signalling is unknown and moreover, is tissue specific. Thus, in the notum, *sca* mutants have an excess of bristles, a phenocopy of *N* mutants, indicating that Sca positively modulates N-activity ^12^. In wing discs, ectopic expression of *sca* inhibits N-pathway by reducing the interactions with its ligands ^13^. In eye discs, some studies indicate that Sca promotes N activation in response to Dl ^14^ and others show that expression of *sca* in receptor precursor cells reduces N-activity ^15,16^.

In this work, we investigated how resumption of SOP division within the notum influences neurogenesis and ultimately fly behavior. In a kinetic study, we show that SOP cells enter mitosis successively according to a temporal wave that propagates from the center towards the anterior and the posterior border of the thorax. We took a genetic approach to demonstrate that the Notch-ligand Delta and the secreted Notch-signaling modulator Scabrous control this wave. By altering cellular morphology, we show that propagation of the wave is mediated by cell-cell interactions through cell protrusions from SOPs. Finally, we present evidence showing that this mitotic wave has an impact on neural wiring and cleaning behavior. Overall, our data support the idea that timing of neural progenitor divisions controls normal wiring in the central nervous system.

## RESULTS

### SOP mitosis resumes in a temporal wave

To monitor mitosis in SOPs, transgenic pupae at 15h APF expressing GFP under the SOP specific *neuralized* driver (*neurD>GFP*) were monitored with live imaging. As a way to comparing nota, the first cell in each row to divide was used as both a temporal (division time, min) and a positional reference (rank, μm), designated SOP_o_. Centering our attention on SOPs located in the dorsal most region of the thorax (Rows 1-3 in Fig 1A), we observed that the SOP_o_ cells were located in the medial region of each row and generally, the more distant a SOP was from its corresponding SOP_o_, the later it divided (Fig 1A-B, S1A and Movie 1). The order of mitosis of SOPs was not strict. However, plotting the time of SOP division against its rank relative to SOP_o_, reveals that on average SOPs resumed the cell cycle in a wave of mitoses spreading towards the anterior and the posterior ends along each row. Thus, all SOPs in a row divided in two hours for all rows analyzed (Fig 1B-C). No statistical difference was found between the rate of the mitotic waves in rows 1, 2 and 3 (*p* = 0.3954, ANOVA) or between left and right rows (*p* = 0.3038, ANOVA). SOP_o_ were not located precisely in the middle of each row, so the number of SOPs in the anterior and posterior portion of each row was unequal (Fig S1–S2). Still, no statistical difference was found between the rates of the anterior and posterior mitotic waves (*p* = 0.0689, ANOVA).

**Figure 1.**
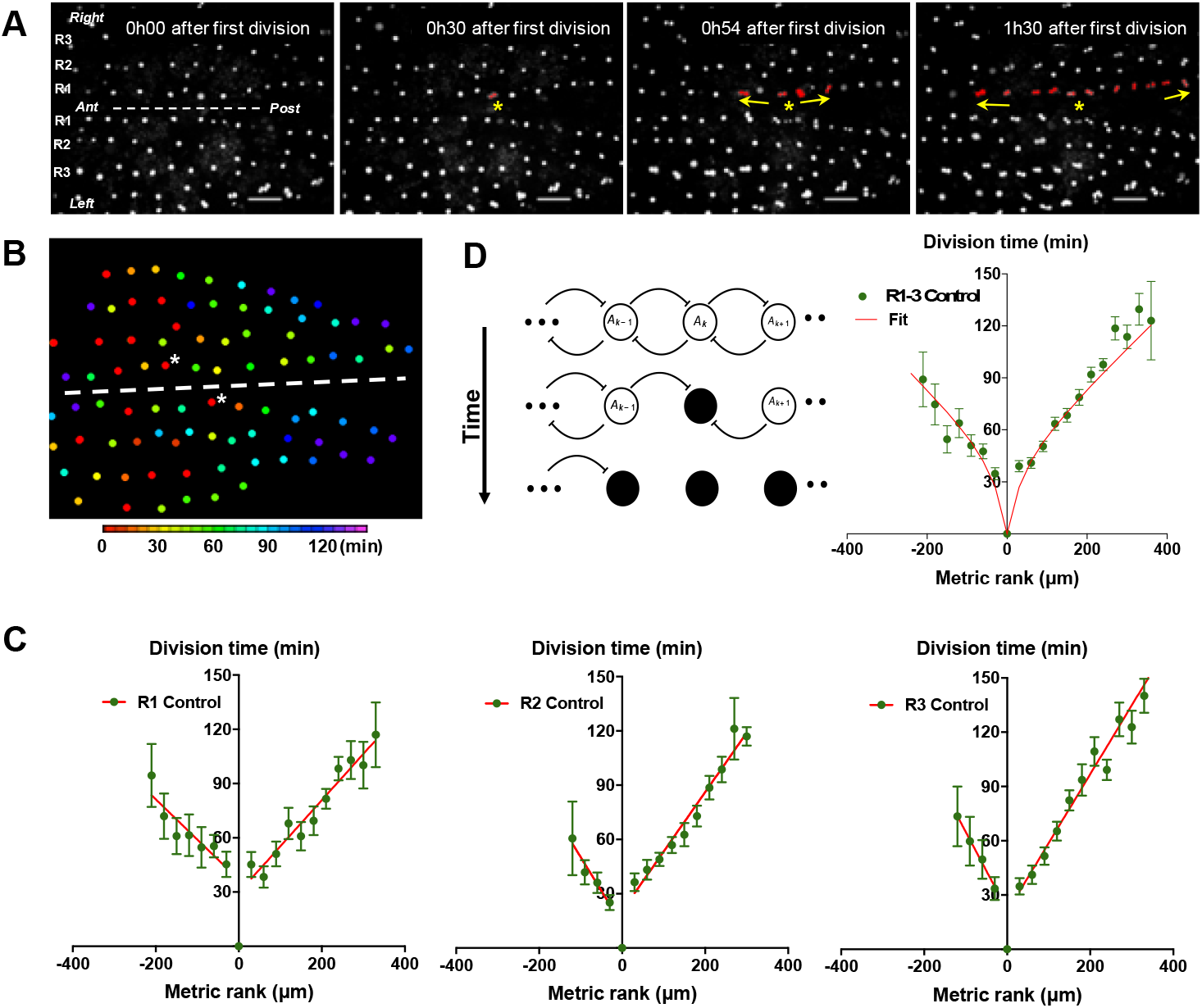
G2-arrested SOP cells of the neuroepithelium resume division in a temporal wave. (A) Four frames of a time-lapse recording of a control *Drosophila melanogaster* pupa from Movie 1. The first three rows of cells in the most dorsal part of the thorax are indicated as R1, R2 and R3 respectively. The dashed line marks the midline, with anterior (Ant) to the left and posterior (Post) to the right. SOPs that have divided are highlighted in red in R1. The yellow arrows indicate the propagation of the wave from SOP_0_ marked by a yellow star. Scale bar, 50μm. (B) Heatmap of a representative control notum depicting time of SOP division. Circles represent SOP cells arranged in rows. The relative time of division is colour coded according to the scale, each colour covering 6 minutes. Only rows 1 to 4 are represented. The dashed line shows the midline, with anterior to the left, and SOP_0_ in row 1 marked by white stars. (C) The time of SOP cell division (mean time ± SEM of n = 16 nota) in row 1 (R1), row 2 (R2) and row 3 (R3) is plotted according to position both relative to the position and time of division of the first cell to divide in each row. Negative and positive metric rank therefore corresponds to SOPs that are anterior and posterior to the SOP_0_ respectively. Red lines show standard linear regressions. SOP cells were identified by GFP expression in a *neur>GFP* fly line. (D) Modelling the wave of SOP divisions. Left, a schematic view of the model at three consecutive times. Empty circles depict non-dividing cells. The flat-headed arrows indicate the inhibition that one cell exerts on immediate neighbours preventing entry into mitosis. Filled circles represent mitotic or post-mitotic cells which no longer have an inhibitory effect thus allowing neighbours to divide. These interactions allow the progression of mitosis along a row. Right, fit of experimental data with theoretical values obtained with the model using as parameters *ρ* = 20 and *μ* = 0.65 (See Model description).

To characterize each curve, the rate of the mitotic wave was calculated as the mean absolute value of the inverse of the slopes of the linear approximation of the curves for all conditions. For control conditions, this gives for the mean slope 3.14 μm min^−1^ (see Table S1). We conclude that the resumption of SOP mitosis is not random because the first division is in a specific region of the notum and a steady wave of division propagates towards the anterior and the posterior parts of the notum.

To establish working hypotheses on the mechanism involved in the propagation of this mitotic wave, we developed a simple contagion model assuming that (1) neighboring progenitor cells in a row inhibit each other from entering mitosis through cell-cell contact, (2) this inhibition is switched off when cells divide, (3) mitoses occur whenever a cellular compound concentration reaches a certain threshold and (4) this compound is produced at a constant rate.

Given these assumptions, the system can be modelled by the differential equations:

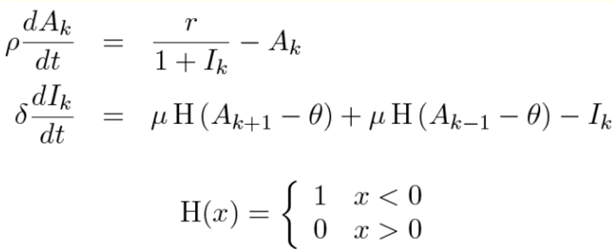

Where *k* indexes cell position. *A* describes the concentration of a pro-mitotic factor with a constant production rate r. Cells divide whenever A is above a threshold *θ*. *I* describes the concentration of an inhibitory compound that inhibits *A*’s production rate. *I* is also constantly produced with a rate *μ* in non-divided neighbouring cells (when A < *θ*) and is not produced in divided cells (A> *θ*). *ρ* and *δ* are degradations times of *A* and *I* respectively. With this set of differential equations - two for each cell - our model accounts for the observed propagation of SOP mitoses very well (Fig 1D).

In particular, this simple model accounts for the non-linearity of the curve observed. Indeed, the interplay between the progressive capacity of SOP cells to divide (modelled by the factor A) and the inhibitory effect on mitotic entry due to neighboring cells (modelled by the factor I that inhibits A’s production) brings cells farther from the start closer and closer to threshold to divide. The result is that, proportionally, cells take less and less time as the mitotic wave progresses along the row. Therefore, the mitotic wave accelerates over time. In other words, the relation between division time and SOP position is flattened for SOP position values far from the origin. This flattening is particularly observed for low inhibition conditions (Fig 1 in Model description). Conversely, at a very strong level of inhibition, the resulting curve is almost linear with a slope different to zero (Fig 1 in model description). The non-linearity of the curve is accentuated between the 1st and 2nd division by the fact that the point 0, the location of SOP_0_, is also the temporal origin. As such, the time of division of SOP_0_ is for definition 0 and it is not a mean as the other points. Since all other values are positive (all cells divide obviously after the SOP_0_), this induces a positive jump in the delay between the 1st and following divisions.

Using this model, variations of only two parameters, *ρ* and *μ*, account for our experimental data (See Parameters estimation in Model description). In the control condition, the best fit was obtained with values of *ρ* and *μ* of 40.38 and 0.52 respectively (Table S2).

### Cytoplasmic protrusions are required for the SOP mitotic wave progression

Based on this model, SOPs are likely to influence neighboring SOPs, so we analyzed the potential ways by which they interact. SOPs in each row are actually separated by 3-4 epithelial cells, but SOPs produce dynamic actin-based protrusions several cell diameters in length that physically interact with protrusions from neighboring SOPs and epithelial cells (Fig 2A left) ^12,17,18^. When SOPs divide, they become spherical and protrusions are drastically reduced (Movie 2). To test the possibility that these protrusions control the mitotic wave, we overexpressed a dominant negative form of Rac1 (*rac1^N17^*) (Fig 2A right and Movie 3), because Rac1 is required for the growth of protrusions in SOPs ^17^. To estimate the protrusion length, we have measured the protrusion extension area (Fig 2B). Under these conditions, SOP protrusions were shorter than in the control (Fig 2C, *p* < 0.0001, ANOVA). Although SOP divisions started at similar developmental times (control 15.7 ± 0.5 h APF, n = 9; *rac1^N17^* overexpression 15.9 ± 0.5 h APF, n = 6), we observed that SOPs with shorter protrusions divide more synchronously than in the control (Fig 2E, mitotic wave rate 6.95 μm min^−1^). Indeed, all SOPs in a row divided within one hour for all rows analyzed (Fig 2E, Row 1 *p* < 0.0001; Row 2 *p* = 0.0292, Row 3 *p* = 0.0014, ANOVA, and Fig S3B). The lesser slopes of the time-rank curves of the mitotic wave in flies with short SOP protrusions shows that neighboring SOPs resume mitosis earlier than normal. These data indicate that SOPs exchange inhibitory signals throughout protrusions to maintain neighboring SOPs in G2-arrest. When a SOP divides, its protrusions are retracted so the inhibitory signal is suppressed allowing progression of the mitotic wave to the next SOP along the row. Interestingly, after fitting the experimental data with the model, the best fit was obtained with no change in the ρ value (in row 1, 40,38 in control and 39,12 in *rac1^N17^*) whereas the value of the inhibitory parameter (μ) decreases (in row 1, 0,52 in control and 0,36 in *rac1^N17^*). For the other rows, the parameter values for the best fit are shown in Fig S4B and Table S2.

**Figure 2.**
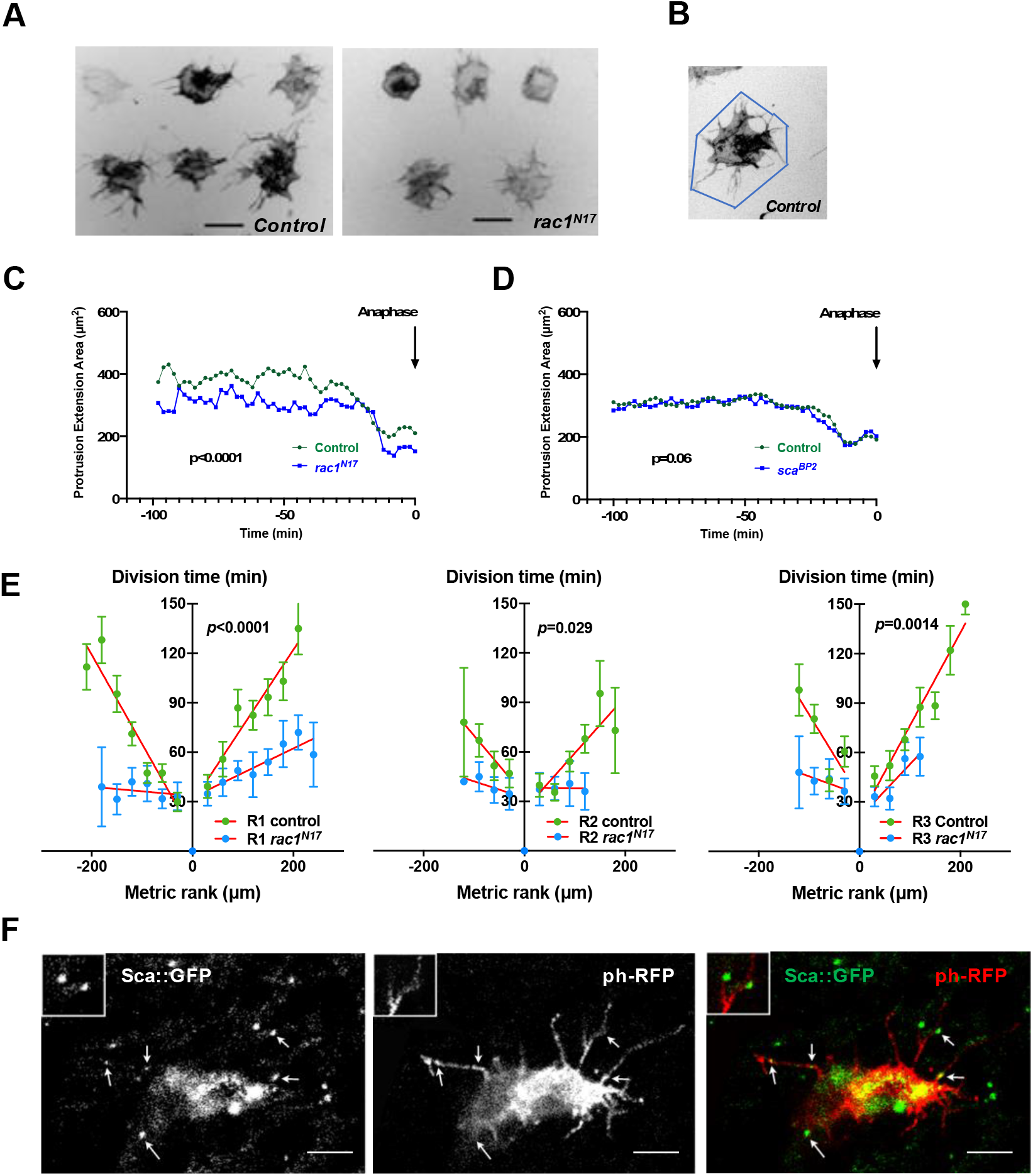
The SOP mitotic wave is controlled by signals exchanged via cell protrusions. (A) Frames from live recordings showing cell protrusions of SOPs in control (left) and *rac1^N17^* (right) pupae. Protrusions were visualized by specific expression of a membrane-tethered RFP form (*ph-RFP*), invert fluorescence. Scale bar, 20μm. (B) Illustration of the procedure to quantify the cell protrusion length. (C) Protrusion extension area over time in control and after *rac1^N17^* overexpression (mean ± SEM, n = 11 cells for control and n=14 for *rac1^N17^*) measured from live recordings. Anaphase was taken as a temporal reference. (D) Protrusion extension area over time in control and in *sca^BP2^* null mutant (mean ± SEM, n = 20 cells for control and n=18 for *sca^BP2^*) measured from live recordings. Anaphase was taken as a temporal reference. (E) The SOP mitotic wave in row 1 (R1), row 2 (R2) and row 3 (R3) in control and after overexpression of *rac1^N17^*. Mean division time ± SEM of n = 6 nota. (F) Sca localisation in a SOP cell. Arrows indicate distinct Sca foci along cell protrusions. Scabrous (left panel) in SOP cell marked with a membrane bound ph-RFP (middle panel), merged in right panel. Images are a maximum projection of three confocal sections. Two Sca foci are magnified in the inserts. Scale bar, 5μm.

### The SOP mitotic wave is regulated by Scabrous and Delta

Cellular protrusions allow molecules to be exchanged between cells for direct and selective signaling ^19,20^. The candidate for such a signal would be a secreted molecule with a role in regulating the interactions and spacing of cells in the nervous system. One such molecule is the fibrinogen-like protein Scabrous (Sca) that regulates several processes such as the regularly spaced pattern of ommatidia in the eye or of bristles in the notum ^12,17,21,22^, or ommatidial rotation ^23^. In the protein trap line *sca::GFP*, Sca was detected as multiple spots of fluorescence in the cytoplasm and along the protrusions of SOPs (Fig 2F and movie 4). At 14h30 APF, very faint Sca spots were detected on the notum. The intensity of these spots reaches a peak at 24h APF then decreases rapidly and the spots were undetectable at 32h APF (Fig S5). To investigate whether Sca is involved in the synchronization of SOP mitosis, we studied the mitotic wave in the *sca^BP2^* null mutant, which is viable at the developmental period studied. In this mutant, we observed SOP_o_ cells at different points along each row as they were not limited to the anterior region (Fig 3A, S1B and S2B-C). More importantly, in the *sca^BP2^* mutant, SOPs divided more simultaneously than in the control (Row 1, *p* = 0.0015; Row 2, *p* = 0.0029; Row 3, *p* = 0.0014, ANOVA, Fig 3A-B, Figs S1, S6A, Movie 5). For instance, Row 1 in *sca^BP2^* divided in one-third of time taken for the control SOP row to divide, and the rate of the wave increased to 20.27 μm min^1^ (Table S1). The release of the inhibition observed in *sca^BP2^* context was again well confirmed by fitting the experimental data with the model. For instance, for row 1, the best fit was obtained with no change in the ρ value (37,71 in control and 41,08 in *sca^BP2^*) whereas the value of the inhibitory parameter (μ) decreases from 0,52 (control) to 0,31 (*sca^BP2^*). The invariance of the ρ and the reduction of μ values were observed for all rows (Fig 3C, Fig S4 E-F and Table S2).

**Figure 3.**
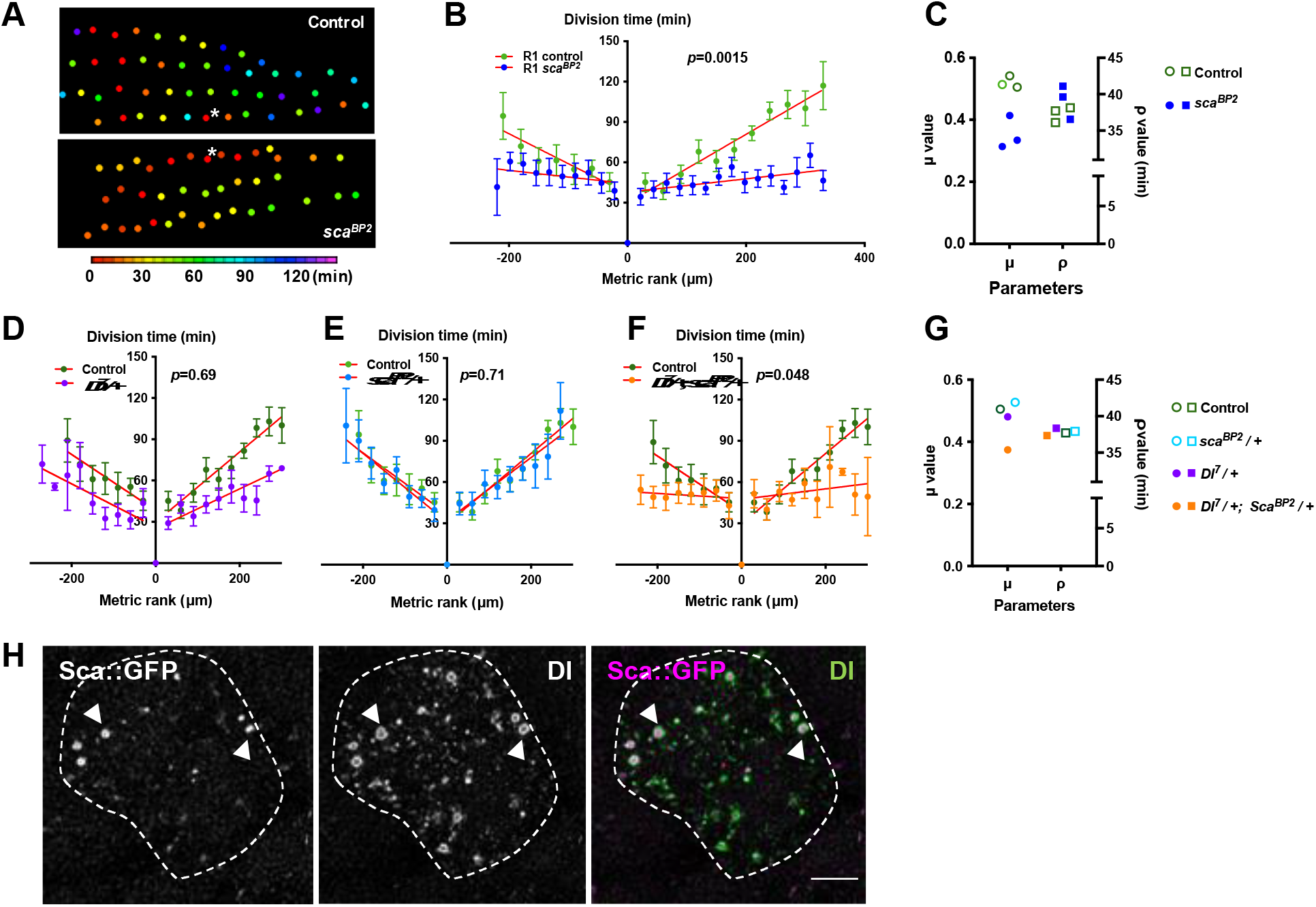
The mitotic wave of SOP cells is controlled by Notch/Delta/Scabrous signals. (A) Heatmap of a representative heminota in control and *sca^BP2^* homozygous null mutant aligned, anterior to the left. The relative time of SOPs division is colour coded according to the scale. (B) The SOP mitotic wave in row 1 in control and in *sca^BP2^* homozygous mutant. (C) Dot plot showing the distribution of μ and ρ parameters extracted from the best fits displayed in Fig S4 D, E. Note that the best fit was obtained in *sca^BP2^* null mutant after reduction of the μ parameter only. (D-F) Genetic interactions between *Dl^7^* and *sca^BP2^*. Comparison of the SOP mitotic wave in control and *Dl^7^/+* (D), *sca^BP2^/+* (E) and *Dl^7^/+, sca^BP2^/+* (F). Mean division time ± SEM, n = 9, 8 and 11 nota respectively. (G) Dot plot showing the distribution of μ and ρ parameters extracted from the fits displayed in Fig S7. Note that in simple heterozygous backgrounds, μ and ρ were similar to the control while in *Dl^7^/+, sca^BP2^/+* double heterozygous background, the best fit was obtained after reduction of the μ parameter only. (H) Confocal image of one SOP in a *sca::GFSTF* protein-trap fly at the moment of the SOP wave (16 h APF), immunostained for GFP (left panel) and Dl (middle panel), merged in the right panel (purple and green respectively). Arrowheads show co-localization of Dl and Sca into vesicle-like structures with Sca::GFSTF as puncta surrounded by Dl staining. SOP is delimitated by a dashed line. Scale bar, 5 μm.

This Sca mediated inhibition is not likely to be related to protrusion length shortening because no difference was found between *sca^BP2^* and control SOP protrusions Fig 2D and ^12^. To exclude the possibility that the absence of Sca during the entirely development of *sca^BP2^* mutant could secondarily impact the mitotic wave, *sca* expression was specifically downregulated using a conditional RNAi strategy between 12 and 19 h APF (Fig S6B), that is from just before and during the mitotic wave. In these conditions, as in null mutant, we again observed that the mitotic wave was flattened (Fig S3C; S6C; Row 1, *p* < 0.0001; Row 2, *p* = 0.69; Row 3, *p* = 0.003, ANOVA). Again, this effect was best fitted by a reduction on the inhibitory parameter compared to the control (Fig S4A, C and F and Table S2). These data indicate that Sca transported through cell protrusions specifically controls the wave of SOP mitoses along the rows.

Sca is known to modulate the Delta/Notch (Dl/N) pathway ^24^, so we tested whether this pathway is involved in regulating the SOP mitotic wave. Since null *Dl* mutants are lethal, we studied the mitotic wave in *Dl^7^/+* heterozygous background. We did not observe a statistically significant reduction in the mitotic wave rate in *Dl^7^/+* SOPs (Fig 3D, *p* = 0.69, ANOVA). Similarly, we did not observe any difference in the rate of the SOP division wave in the *sca^BP2^*/+ heterozygous line (Fig 3E, *p* = 0.71, ANOVA). By contrast, when the gene dosage of both *sca* and *Dl* was reduced by half, the time-rank curves were flattened corresponding to a significant increase in the mitotic wave rate (from 3.14 to 43.35 μm min^-^1) (Fig 3F, *p* = 0.048, ANOVA). Again, this effect was best fitted by a reduction on the inhibitory parameter (Fig 3G, S7 and Table S2). These results demonstrate that *sca* and *Dl* genetically interact, which suggests that these factors act in concert to control the SOP mitotic wave. Indeed, we observed that Dl and Sca co-localize in vesicle-like structures inside the cytoplasm with Dl surrounding the Sca staining (Fig 3H). This co-localization firstly described Renaud and Simpson (2001)^12^, evidences that these two proteins are transported from the endoplasmic reticulum to the membrane in the same vesicles.

Altogether, our results show that the wave of SOP mitoses along rows depends on an inhibitory signal transmitted by protrusions from the progenitors and is controlled by Dl and Sca proteins.

### Outcome of the mitotic wave on microchæte axonogenesis

Considering the invariability of sensory organ formation, we wondered whether differences in the wave of SOP mitosis might impact the timing of axonogenesis along each row. To this end, we measured axon length at the beginning of axonogenesis (24 h APF) in control and *sca^BP2^* pupae (Fig 4A). In control pupae, the axon length varied along the antero-posterior axis. Indeed, within a row, more the neurons were located at the extremities, the shorter their own axon. However, in *sca^BP2^* animals, the length of the axon along a row was significantly more homogeneous (Fig 4B, *p* = 0.0232, ANOVA). These observations indicate that as a result of the SOP mitotic wave, neurogenesis and consequently axonogenesis progress from the centrally located neurons towards those located at the extremities in controls, while in *sca* mutants, in which the wave was either reduced or absent, axonogenesis occurs simultaneously along each row. Thus, the mitotic wave of SOP division controls axonogenesis timing.

**Figure 4.**
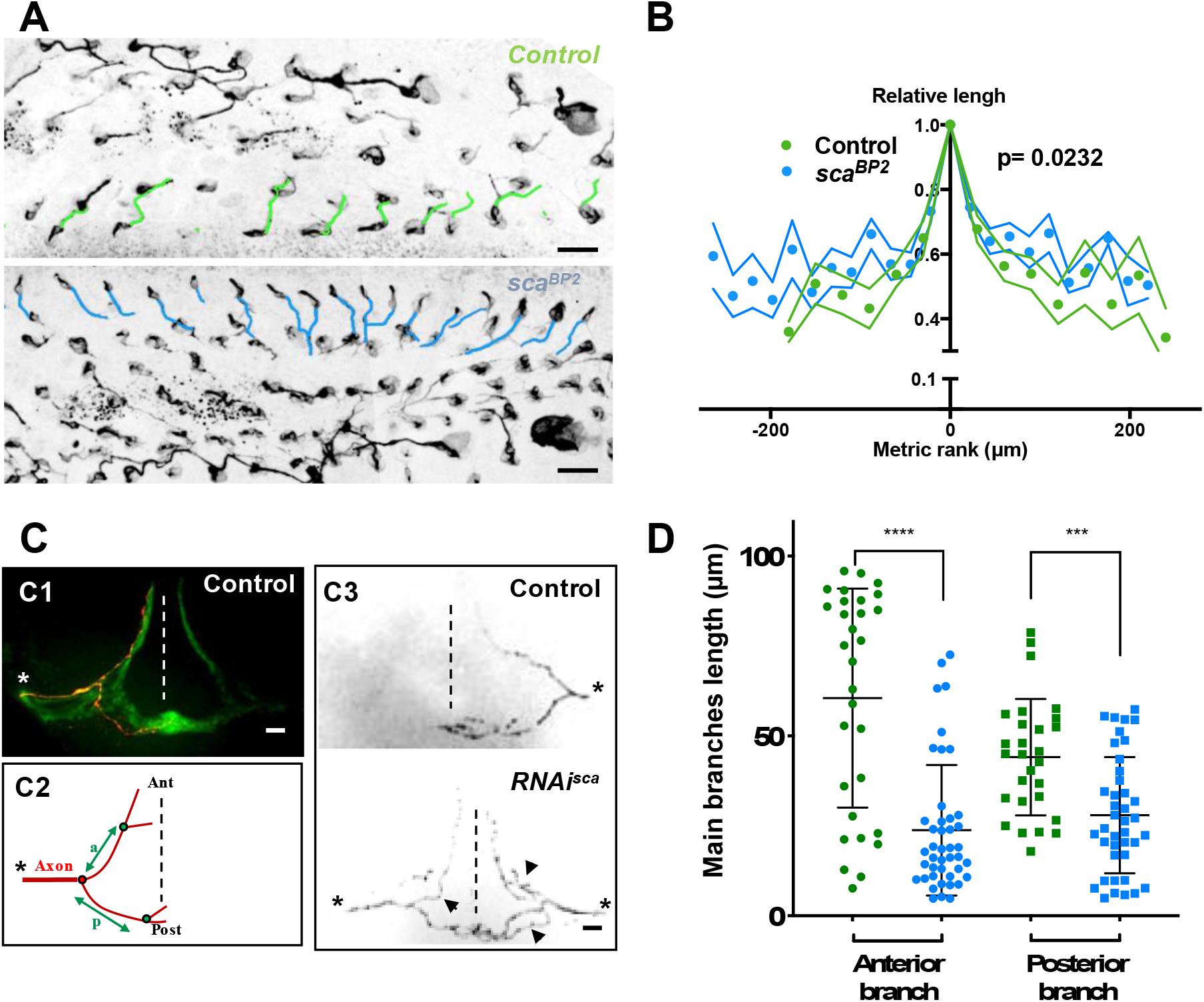
Microchæte axonal projections are affected by the amplitude of the SOP mitotic wave. (A) Axon immunostaining using anti-Futsch antibodies in control (*w^1118^*) and *sca^BP2^* heminota at 24h APF aligned, anterior to the left. Axons in row 1 were artificially coloured in green for *w^1118^* and blue for *sca^BP2^* homozygous flies. (B) Relative axon length of neurons belonging to sensory organs on row 1 plotted against relative position (mean length ± SEM of n = 18 rows). The neuron with the longest axon on the row was taken as temporal and spatial reference in control (*w^1118^*) and *sca^BP2^* homozygous pupae at 24h APF. (C) Axon projections of single sensory organ neurons in the thoracic ganglion identified with MCFO strategy. The midline is indicated by vertical dotted lines, anterior (Ant) up and posterior (Post) down. The asterisks indicate the point of entry of the sensory nerve into the thoracic ganglion. (C1) A single axon projection (red) counterstained with the neuropile formed by all axon terminals from sensory organs on the central notum (green). (C2) Schematic drawing of an axon projection. The length of the anterior (a) and posterior (p) primary branches were used to quantify the degree of branching of axon terminals. (C3) Representative examples of axon projections into the thoracic ganglion in control fly (top panel) and when the SOP mitotic wave was abolished (*RNAi-sca*) (bottom panel). Arrowheads point to additional branches in *RNAi-sca*. (D) Lengths of anterior and posterior primary branches in single axon terminals of control (green, n = 30) and in flies where the SOP mitotic wave was abolished (*RNAi-sca*) (blue, n = 43). Mean ± SEM \*\*\*\**p* < 0.0001, \*\*\**p* = 0.0001, two-tailed unpaired t-test.

### Loss of the SOP mitotic wave leads to a modified organization of microchæte axonal projections

We wondered whether the altered axonogenesis timing observed in the mutant bristle organs influences the patterning of axonal endings in the thoracic ganglion. To address this question, the morphology of these axon terminals was studied by expressing a membrane-bound GFP form in bristle cells using pannier-GAL4, as a driver of dorsal expression. Under these conditions, the neuropile in the thoracic ganglion formed by axon terminals from all bristles of the central dorsal notum was visualized as a mesh-like triangular structure formed by an anterior arm at each left and right sides and a posterior commissure that cross the midline (Fig 4C1). In 11% of ganglia analyzed in control flies a second medial commissure was observed (Fig S8 arrow). Interestingly, in flies in which the mitotic wave was abolished after *sca* downregulation (*RNAi-sca*), this percentage increased to 34% (Fig S8, *p* = 0.0002, Fisher’s exact test). These observations prompted us to analyze single axon terminals by immunodetecting individual axons using the stochastic labelling method MCFO ^25^. We observed in control flies that individual axon terminals have two main primary branches, an anterior branch that does not cross the midline and a posterior branch that crosses the midline (Fig 4C1 and C3 top panel). In *RNAi-sca* animals, we observed that axon terminals are more branched (Fig 4C3 bottom panel). The lengths of primary anterior and posterior branches, between the root and the first branch point (Fig 4C2), of *RNAi-sca* axons were significantly shorter than those of control axons (Fig 4D, *p* < 0.0001 and *p* = 0.0001 respectively, ANOVA). Since on the one hand, *sca* was inactivated specifically and only during the mitotic wave, and on the other hand, *sca* expression turn-off prior to axonogenesis (Fig S5), the effects observed could not be due to a possible effect of *sca* on axonogenesis itself. As primary branches were shorter when SOP division occurred simultaneously on the notum, the timing of axonogenesis is likely to be an important factor controlling axon branching. Interestingly, our data are in agreement with studies showing that axon terminals of bristles from rows that are formed earlier, are more branched than those from rows that develop later ^9^. As such, our results suggest that the order of birth of neurons in a given row contributes to the future connectivity of these neurons.

### Loss of the SOP mitotic wave leads to changes in fly behavior

Finally, we wondered whether modifications in axon branching after disruption of normal timing of SOP mitosis led to behavioral changes. In order to test this possibility, we analyzed the cleaning reflex, as a read out for the function of the mechanosensory system of flies, in conditions where the SOP mitotic wave was abolished. The cleaning reflex is a patterned set of leg movements elicited in a fly when its thoracic bristles receive tactile stimulation (Fig 5A and Movie 6) ^10,26^. When the SOP mitotic wave was impaired, after specific donwregulation of sca during the wave, we found that the number of air puffs required to elicit a leg response was significantly higher than in control flies (Fig 5B, *p* = 0.028, ANOVA).

**Figure 5.**
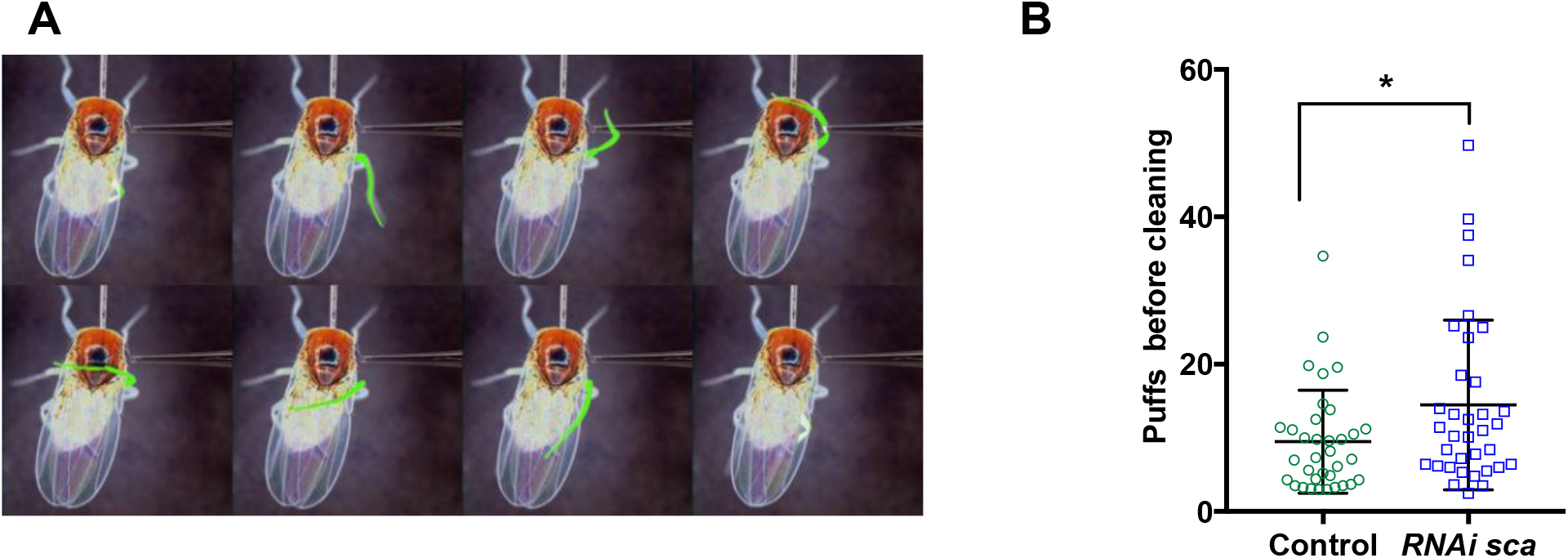
Loss of the SOP mitotic wave leads to changes in fly behavior. (A) Eight frames of a time-lapse recording of a decapitated control fly showing the sequence of movements upon stimulation of the dorsal bristles. (B) Number of air puffs required to elicit a cleaning reflex in control flies (n = 35) and in flies where the SOP wave was abolished (*RNAi-sca*) (n = 35). Mean ± SEM. \**p* = 0.0238; ANOVA.

All together, our results show that the temporal order of neural precursor cells division impact on neuronal cell diversity and ultimately on behavior.

## DISCUSSION

To study how functional neuronal diversity can be generated from a homogenous set of neural precursors, we took advantage of the invariant way in which sensory organs are located on the dorsal epithelium of *Drosophila*. This spatial configuration greatly facilitated the study of the relative timing of SOP division and the identification of a distinct temporal wave of SOP mitosis. Asynchrony in mitotic reactivation timing has been described in *Drosophila* larva neuroblasts. This differential timing is related to two cell cycle arrests: one population of neuroblasts is arrested in G2 while another population is arrested in G0 ^27^. G2-arrested neuroblasts resume mitosis earlier than those in G0-arrest. As in our system, it has been proposed that this particular order of division ensures that neurons form appropriate functional wiring. It is relevant that other temporal processes controlling the wiring of peripheral receptors with the central nervous system have been described in the *Drosophila* eye, another highly organized structure ^28^. It is conceivable that these temporal patterning mechanisms of neurogenesis, to date identified only in organized tissues, could be more widespread.

A core aspect of our work was to link cellular level of complexity (timing of SOP division) with uppermost level (behavior). In this context, we presented evidences showing that the cleaning reflex was impaired when the SOP mitotic wave was abolished. The cleaning reflex has been traditionally analyzed after stimulation of macrochaetes rather than microchaetes as in the present work. Macro- and microchaetes have different patterns of terminal axon arborization ^9^. As such, it is remarkable that this fly behavior was significantly affected by altering the timing of microchaete precursor division in the dorsal thorax. We show that the SOP mitotic wave leads to a progressive neurogenesis along each row of microchaetes. This, in turn, would likely induce a particular pattern of microchæte axon arrival in the thoracic ganglion required for the proper organization of the neuropila in the central nervous system. Although we have documented this progressive axonogenesis, we do not know the strict pattern of axon arrival into the ventral ganglion. It would depend on the order of birth of neurons, and on the geometry of axon projections that fasciculate to form the dorsal mesothoracic nerves in the ganglion. In any case, we show here that, when *sca* function was specifically downregulated during the SOP mitotic wave, axonogenesis occurs almost simultaneously in each row of microchaetes. This certainly impairs the pattern of axon arrival into the ganglion leading to ectopic axon branching and changes in fly behavior. It would be interesting to know whether these impairments are specifically due to neurogenesis occurring simultaneously. To test this, we need find a way to induce different patterns of SOP mitotic entry, for instance, a centripetal wave or a random order. If the observed effect is specifically due to the simultaneity, normal behavior would be expected to be associated with other patterns of SOP division.

We observed that the first SOP to divide (SOP_0_) was always located in the anteromedial region of each row. This may reflect the existence of a pre-pattern that causes SOPs located in that region to start dividing earlier than the others. Although the anteromedial region corresponds approximately to the posterior limit of expression of the transcription factor BarH1 ^4^, no factors specifically expressed in this region have yet been identified. Alternatively, as the location of SOP_0_ is modified when Sca function was impaired, an interesting possibility is that SOP_0_ is selected by an emergent process related to cell-cell interaction in the epithelium, rather than by a passive pre-pattern that organizes the first events in the notum.

We present evidence indicating that the secreted glycoprotein Scabrous, which is known to interact with the N-pathway to promote neural patterning, controls the kinetics of SOP mitosis in the notum. In proneural clusters, cells that express high levels of Dl and Sca become SOPs, while surrounding epithelial cells activate the N-pathway to prevent acquisition of a neural fate ^30,12,29^. In eye and notum systems, Sca modulates N-activity at a long range. Indeed, during eye development, *sca* is expressed in intermediate clusters in the morphogenic furrow and transported posteriorly in vesicles through cellular protrusions to negatively control ommatidial cluster rotation ^23^. Similarly, in the notum, SOP protrusions extend beyond several adjacent epithelial cells in which Dl and Scabrous are detected by Renaud and Simpson ^12^ and confirmed in this study. Our data show that shorter protrusions (obtained after Rac1^N17^ conditions) as well as loss of function of *Dl* or *sca* make the mitotic wave more synchronous. It is plausible that Sca, transported through protrusions, is required to maintain SOPs in G2 arrest, though we cannot formally rule out the possibility that Rac1^N17^ overexpression affects sca secretion *per se*.

As in neuroblasts, G2 arrest in SOP cells is due to the downregulation of the promitotic factor Cdc25/String. Thus, overexpression of *string* in SOPs induces a premature entry into mitosis ^31^, while overexpression of negative regulators, like Wee1, maintain these cells in arrest ^32^. Possibly Sca negatively regulates *string* expression, perhaps through the N-pathway that it is known to control the level of String ^33,34^. Alternatively, it has been recently shown that the insulin-pathway also regulates String level ^27^. Moreover, in muscle precursors, cell proliferation is induced by the insulin-mediated activation of the N-pathway ^35^. These observations raise the interesting possibility that, in our system, insulin activates the N-pathway and Sca modulates this activation. Further investigations will be required in order to identify the link between Scabrous, the N/Dl- and insulin-pathways in the resumption of mitosis in SOPs.

During nervous system development, the complex patterns of neuronal wiring are achieved through the interaction between neuronal cell surface receptors and their chemoattractive or repulsive ligands present in the environment ^36^. An essential condition for proper axon guidance is the competence of neurons to respond to these environmental clues. It is generally agreed that neuron competence depends on the specific expression of transcriptional factors regulating their identity ^37^. We show here that the timing of neuron formation is also a factor controlling their terminal morphology. We propose that the SOP mitotic wave induces a particular pattern of arrival of microchæte axons in the thoracic ganglion (Fig 6). This pattern establishes a specific framework of guidance cues on which circuits will be built and ultimately influencing an organism’s behavior. Our findings support the idea that, in addition to genetic factors, neurogenic timing is a parameter of development in the mechanisms controlling neural branching.

**Figure 6.**
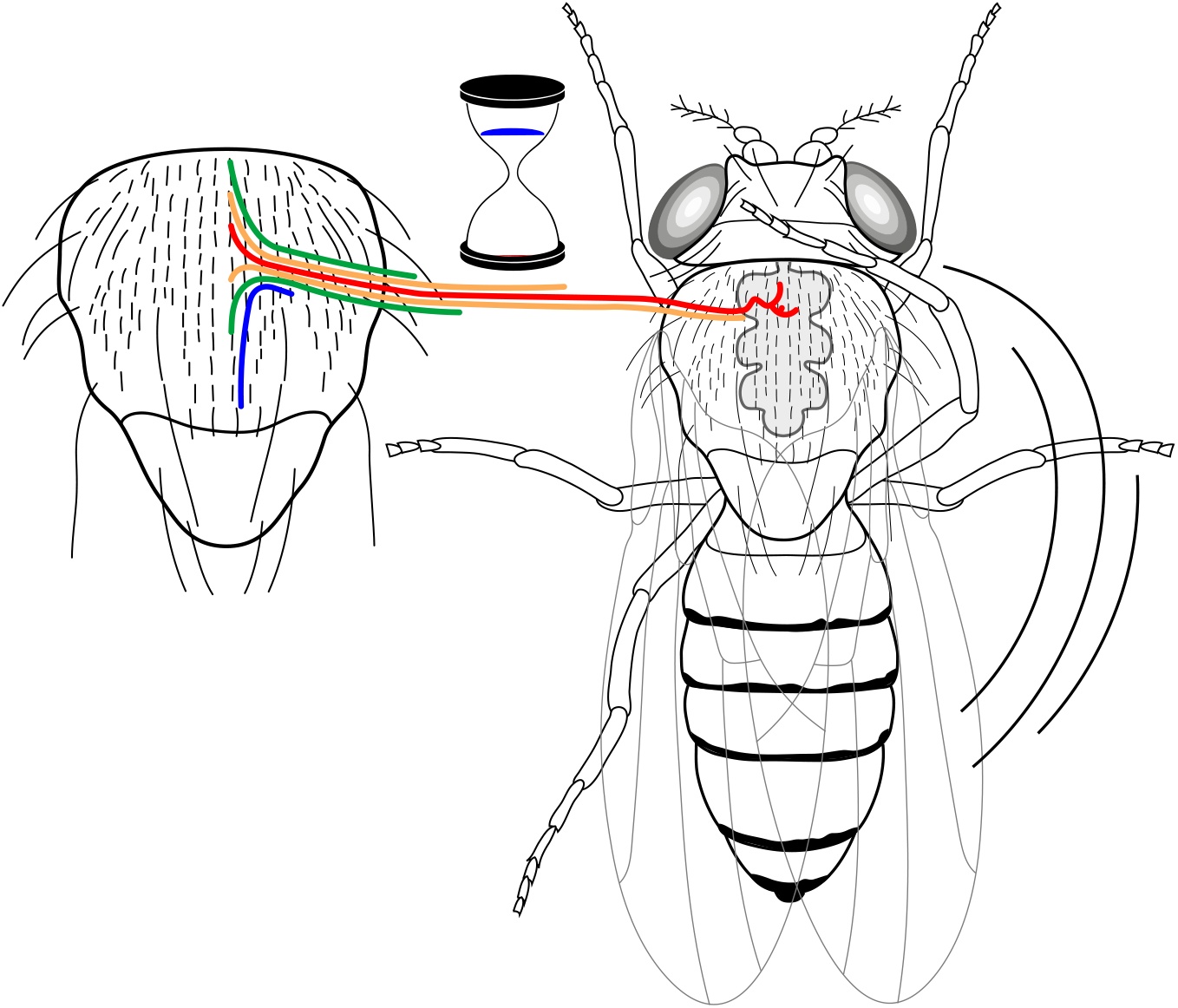
A neural progenitor mitotic wave is required for asynchronous axon outgrowth. The mitotic wave of precursor cells induces an asynchronous arrival of sensory axons into the thoracic ganglion. This is depicted here by axons coloured according to the developmental time of sensory organs that in turn is controlled by the wave of precursor cell divisions. Red, early-formed; blue, late-formed sensory organs. This asynchrony would be associated with a particular axon connectivity in the thoracic central ganglion required for a normal cleaning behaviour in the fly.

## Supporting information

Model Description

## Acknowledgments

We thank Marie-Emilie Terret (CIRB, Paris, France) and Bassem Hassam (ICM, Paris, France) for critical and constructive comments on the manuscript. We thank Sophie Gournet for the Figure 6. We are grateful Julie Perrin and Chloé Jean Baptiste Simonne for their participation in the experiments presented respectively in Figures S8 and 4C-D.

This work was funded by institutional support from the Centre National de la Recherche Scientifique (CNRS) and Sorbonne University.

## Author contributions

Conceptualization: J.L., A.A., M.G., S.L.V.; Methodology: H.S. J.L., M.G., S.L.V.; Formal analysis: J.L., H.S.; Investigation: J.L., A.A., P.D, M.G., S.L.V.; Resources: A.B; Writing Original Draft: J.L., M.G., S.L.V. Funding: M.G.

## METHODS

### Fly strains

Standard methods were used to maintain fly stocks. The *Gal4/UAS* expression system was used to express several constructions in the mechanosensory bristle lineage as listed in the key resources table. Two drivers were used: *neuralized^p72^>Gal4* ^38^ expressed in SOPs and their descendants and *pannier>Gal4* ^39^ expressed in the dorsal most domain all along pupal stage. For temporal control of transgene expression, *Gal4* drivers were combined with *tub-Gal80^ts^*. Crossed flies, developing embryos and larvae were maintained at 18 °C and then pupae were shifted to 30 °C to allow the expression of *Gal4*. For the conditional inactivation of *sca* and the ovexpression of Rac dominant negative form, the shift was done from 12 h APF to 19 h APF at the moment of the mitotic wave. To visualize SOP membranes specifically, we used a transgenic line expressing the pleckstrin homology domain of PLCδ fused to RFP (*ph-RFP*) under the control of the *neur* regulatory sequences (*neur>ph(PLCδ)-RFP* ^40^.

### MultiColor FlipOut (MCFO) labelling

MCFO-1 flies ^25^ were crossed with *UAS-RNAi-sca, pnr>Gal4, tubGal^80ts^*. For the control, the crosses were maintained at 18 °C until the progeny reached adulthood. For conditional inactivation of *sca*, pupae were shifted to 30 °C between 12 h APF to 19 h APF, then returned to 18 °C until they reached adulthood. In both cases, adults were heat-shocked at 37 °C for 12.5 min (to activate flipase expression), then left for two days at 25 °C to allow the expression of tags. After removing the head, the flies were fixed in 4% paraformaldehyde for 48 h and rinsed in T-PBS (0.1% Triton X100 in PBS). After dissection, thoracic ventral ganglions were incubated with primary antibodies overnight at 4 °C. After three T-PBS washes, ganglions were incubated with secondary antibodies for 1h at room temperature. Ganglions were mounted in Glycerol-PBS (80% glycerol in PBS) and imaged the same day.

### Immunostaining

Dissected nota from pupae at 16 h APF for Scabrous detection or 24 h APF for axon labelling were processed as described previously ^7^. The antibodies used are described in the key resources (Table 2). Incubations with primary antibodies were done overnight at 4 °C and with the secondary antibodies for 1h at room temperature. Nota were mounted in Glycerol-PBS (80% glycerol in PBS, 1% propylgalate).

### Confocal microscopy

Immunostaining was observed with an Olympus BX41 fluorescence microscope (objective 40X/1.30 or 63X/1.25) equipped with a Yokogawa spinning disc and a CoolSnapHQ2 camera driven by Metaview software (Universal Imaging).

For co-immunodetection of Scabrous and Delta, images were acquired with a Leica SP8 confocal microscope, with the HC PL APO CS2 93 X/1.30 GLYC objective. We tuned the white light laser (WLL) to 650 nm for the excitation. The detector was a HyD. The pixel size was 0.087 μm and the z step size was 0.332 μm. Deconvolution was done with Huygens software. All images were processed with Fiji software ^41^.

### Time-lapse recording

*In vivo* imaging was carried out as described previously ^7^. The temperature during the recording was controlled using a homemade Peltier device.

### Movie and Image analysis

The division time of SOPs was analyzed for each row with Fiji. The first dividing SOP (SOP_0_) provided the “zero time”. For other SOPs, the timing of division was calculated according to this temporal reference. SOP_0_ also defined the “rank zero”. The rank of neighboring SOPs was incremented by one unit from this spatial reference. In order to compare several nota and take account for the neurogenic effect (the increase in the number of SOPs per row) observed in *sca^BP2^*, we measured the distance between SOPs. In *sca^BP2^*, the mean number of SOPs was 18 ± 3 separated by a distance of 22 ± 7.2 μm, while in other genetic backgrounds, there are 13 ± 1 SOPs separated by a distance of 30 ± 9 μm. For each graph, the metric rank corresponds to the rank multiplied by the mean distance between two SOPs.

### Heatmaps

SOPs, identified using fly lines with the *neuralized* fluorescent construction, in pupae from 15 h APF were tracked by live imaging and the time of their division (identified as the anaphase mitotic figures) recorded. The position of cells at the moment of the division as well the relative time of division, encoded according to a rainbow scale, were used to construct each heatmap. The absolute time of division of the first cell which divides in each row was taken as temporal reference. In the dorsal thorax, no migration occurs, so cells keep the same neighbouring and cell positions were roughly those at the start of the live recordings.

### Measurement of protrusion length

Live recording from *neur>ph(PLCδ)-RFP* pupae at 15 h APF were obtained by combining (max intensity) z-stacks (series of confocal optical sections separated by 1 μm) acquired every 3 min using a 40X oil immersion objective. For each SOP analysed, the protrusion extension area was calculated from the convex hull of the polygonal line enclosing the extremities of the longer protrusions at each time frame using Fiji software.

### Measurement of axon length

The length of axons in the nota and of the axon terminals in the thoracic ganglion was measured using the Simple Neurite Tracer plugin of Fiji software. To compare the axon lengths of all the nota, the lengths were normalized to the longest axon in each row. The position of the neuron in each row was defined in the same as in the analysis of time of division. To estimate the extent of axon terminal branching in the thoracic ganglion, we measured the length of primary branches between the root and the first branch point (see Fig 4C2).

### Cleaning reflex assay

Experiments were performed on headless flies. After decapitation, flies were allowed to recover for 1 h, then pasted at the tip of a Pasteur pipette. The most posterior thoracic bristles in the square formed by the four dorso-central macrochæte and the left dorso-central macrochæte were stimulated by air puffs from an Eppendorf microinjector. Specifically, a 0.1-second puff of 15 hPa was delivered every 0.5 second for a minute. For each fly, the stimulation was done 10 times. The number of puffs required to elicit the first leg movement was counted and the mean for each fly was calculated and plotted.

### Statistics

For the mitotic wave study, all statistical analyzes were performed using statistical software R (r-project.org). To compare division time, Linear Mixed Models (LMM) were used. Normality of residuals and homoscedasticity were first assessed. Then LMM was performed using various variables. When a condition was found to be non-significantly different (e.g. Left or Right side of the notum) data were pooled but fly identification was maintained as a fix factor. For example, formula such as this one: lmer(TIME ~ COND + POS + ROW + (1|ID), data=data) was used to assess the impact of TIME according to POS (position) and COND (genetic condition in this example), corrected for ROW and for fly ID as a fixed factor. This test will yield significant influences of factors corrected by the others as well as *p*-values. Post-hoc analyzes were performed for the factor (e.g. ROW) only when the overall model was significant for this factor. For behavioural statistical tests, LMM was used to consider behavioural repetitions using fly ID as a fixed factor and the repetition number.

Unpaired *t* tests and Fisher’s exact tests were done using Prism 7 software.

## Supplementary figures

**Figure S1.**
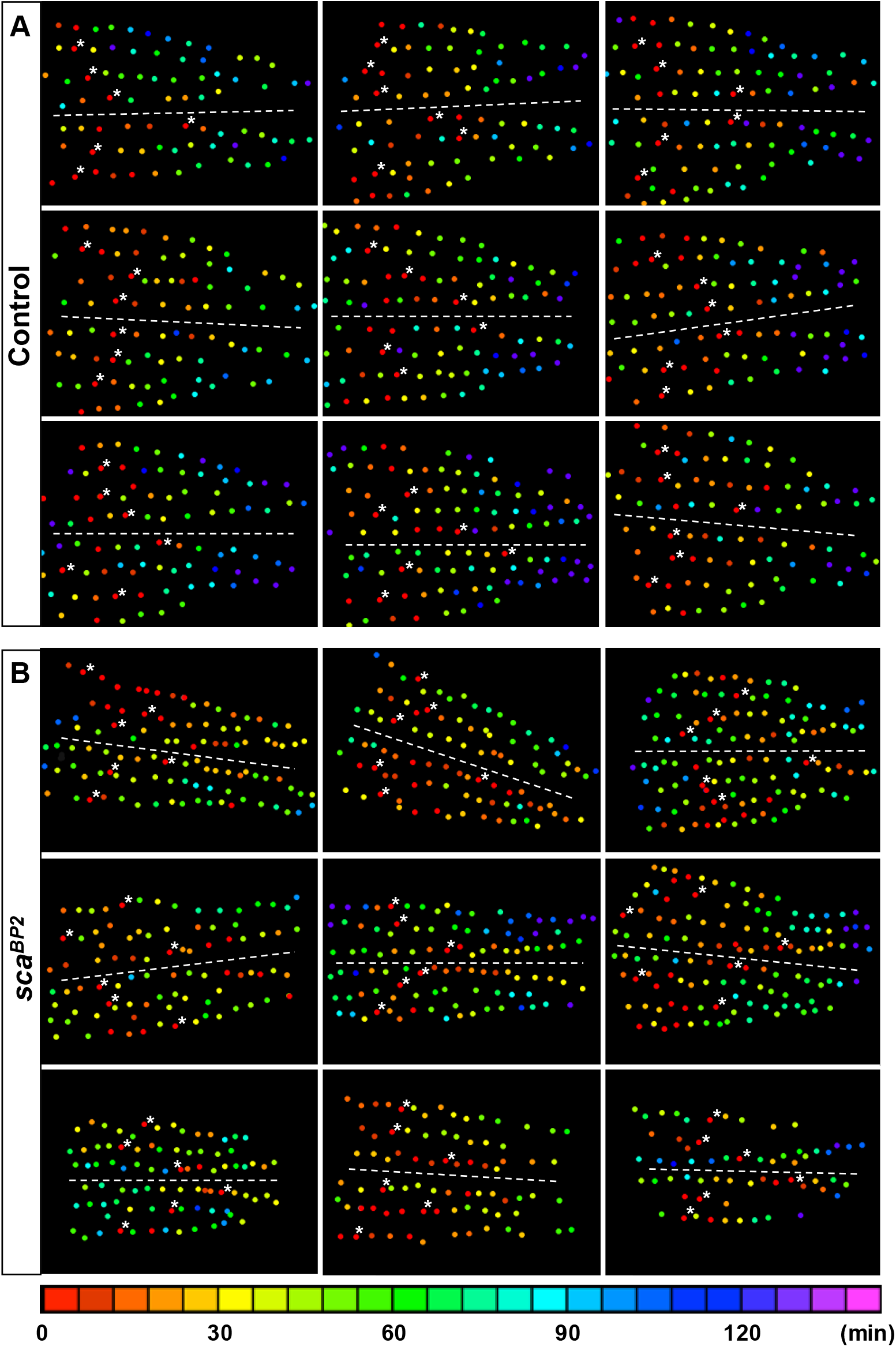
Illustrations of individual mitotic waves in control and *sca^BP2^* mutant. Heatmaps of nine nota showing the time of SOP division in control (A) and *sca^BP2^* pupae (B). Circles represent SOP cells arranged in rows, anterior to the left. The relative time of division is colour coded according to the scale along the bottom, each colour covering 6 minutes. In each row, SOP_0_ are marked by white stars. Dashed lines show the midline. Note that in mutants, many cells are coloured in the reddish part of the spectra showing that several SOP cells divided simultaneously in the same row rather than in control where these cells are located in the anterior region of each row.

**Figure S2.**
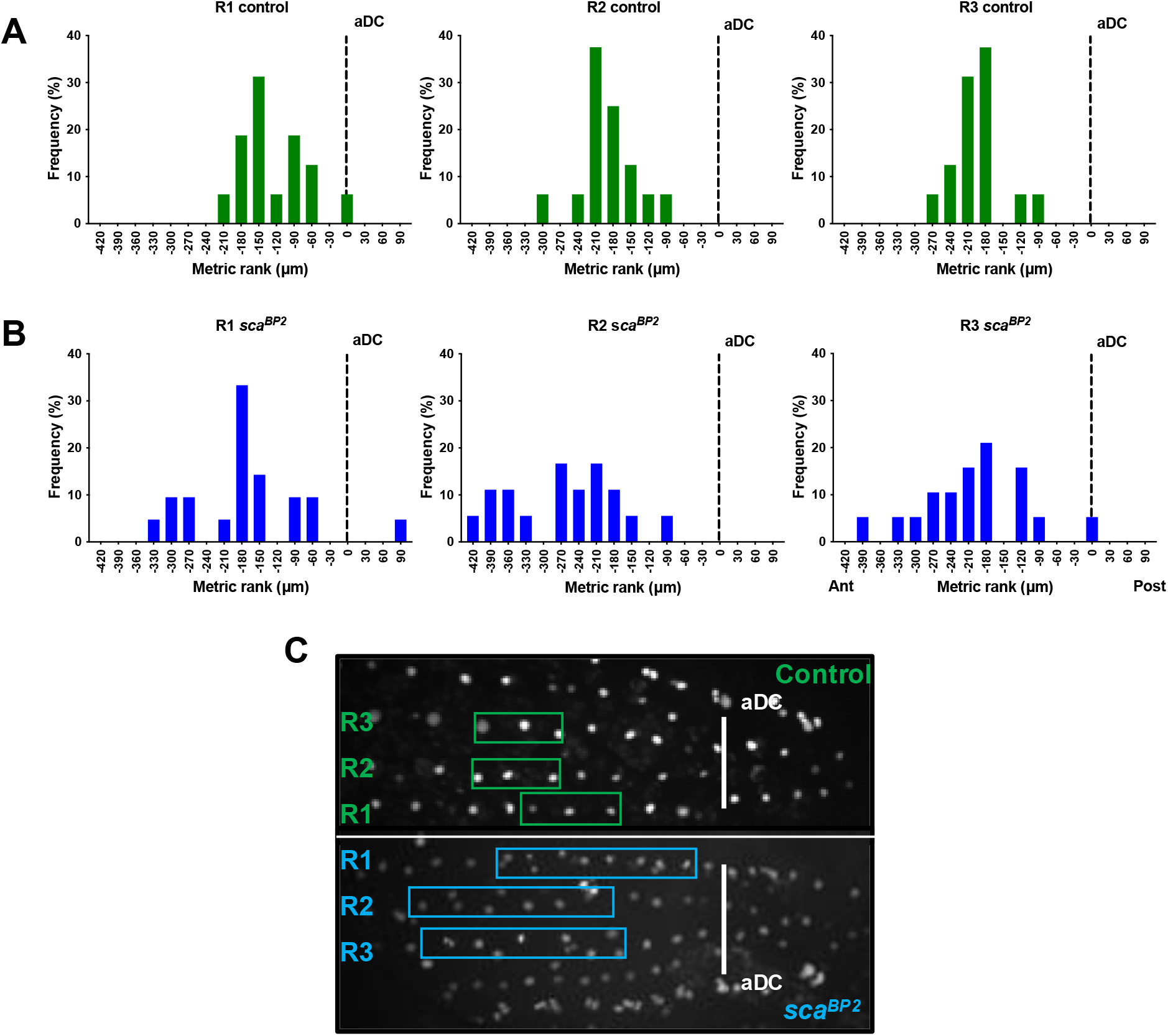
The mitotic wave origin spreads out in *sca* mutant. Location of the first dividing SOP (SOP_0_) in rows 1-3 in control flies (A) and *sca^BP2^* flies (B). The line relaying the two antero-dorsocentral macrochætes (aDC) was used as a spatial reference (dotted line on graphs A and B) (see panel C). Negative and positive metric ranks correspond to SOPs anterior and posterior to the aDC line respectively. Graphs represent the frequency of SOP_0_ at a given position recorded in 9 nota in row 1 (R1), row 2 (R2) and row 3 (R3). (C) Image of hemi-nota of a control fly and a *sca^BP2^* homozygous mutant aligned anterior to the left showing the organisation of the SOPs in the notum. White vertical lines connecting the two aDC were used as spatial reference. Boxes show the locations of 68% of SOP_0_ in each row, corresponding to one standard deviation from the mean.

**Figure S3.**
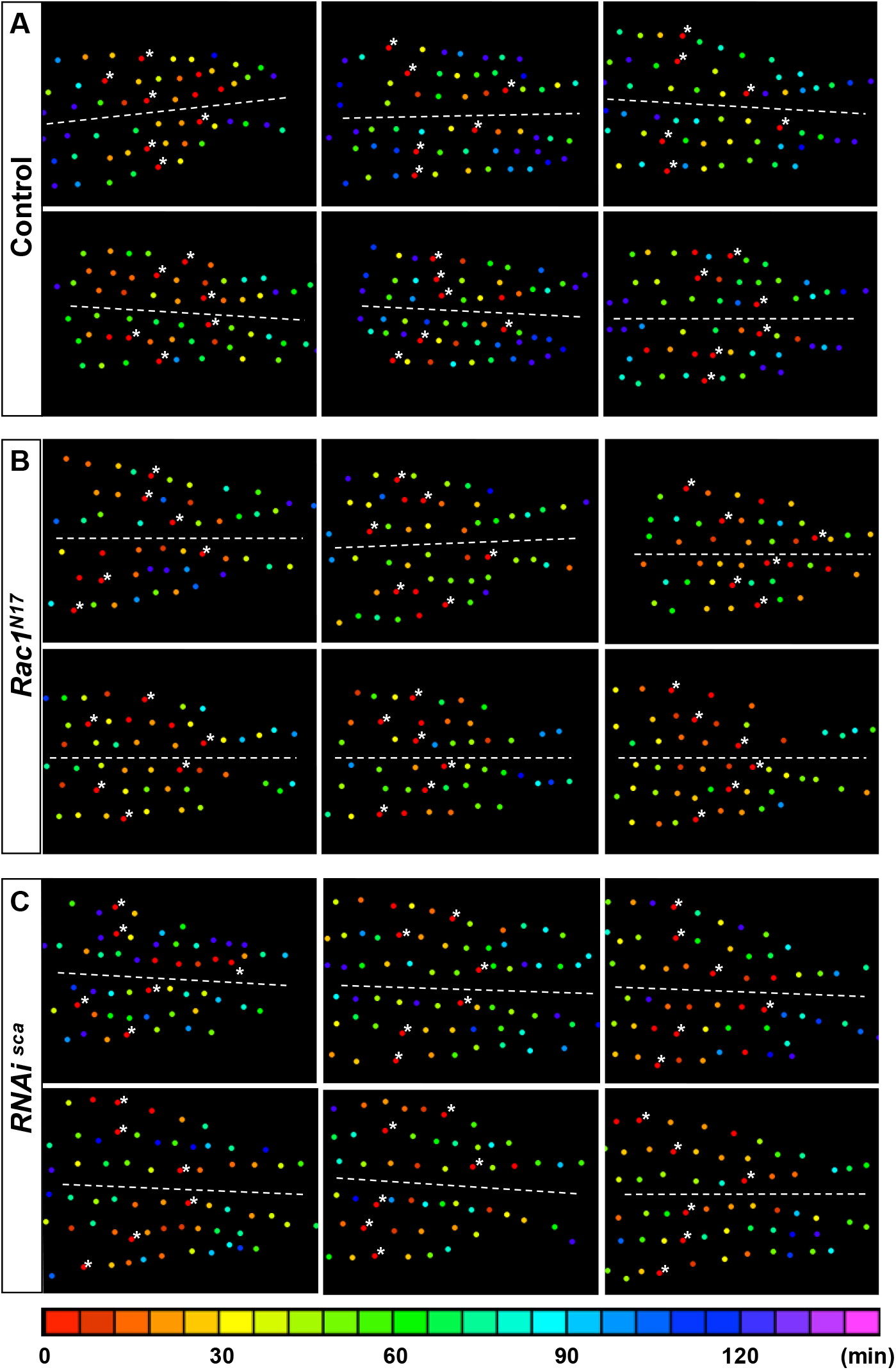
Illustrations of individual mitotic waves after reduction of cell protrusions by overexpression of Rac1^N17^ and after conditional inactivation of *sca*. Heatmaps of six nota showing the time of SOP division in *neur>ph (PLCδ)::RFP/+; pnr>Gal4 tub> Gal80^ts^/+* pupae (A), when cell protrusions were reduced in *neur>ph (PLCδ)::RFP/+; pnr>Gal4 tub> Gal80^ts^/UAS-rac1^N17^* pupae (B) and when *sca* was specifically downregulated between 12 and 19h APF, starting just before and throughout the mitotic wave, using the conditional genetic context *pnr>gal4 tub GAl80^ts^ RNAi-sca* (C). Circles represent SOP cells arranged in rows, anterior to the left. The relative time of division is colour coded according to the scale along the bottom, each colour covering 6 minutes. In each row, SOP_0_ are marked by white stars. Dashed lines show the midline. Note that in (B) and (C) that many cells are coloured in the reddish part of the spectra showing that several SOP cells divided simultaneously in the same row. Note also that the driver *pnr* used to overexpress *Rac1^N17^* disorganizes the pattern of rows.

**Figure S4.**
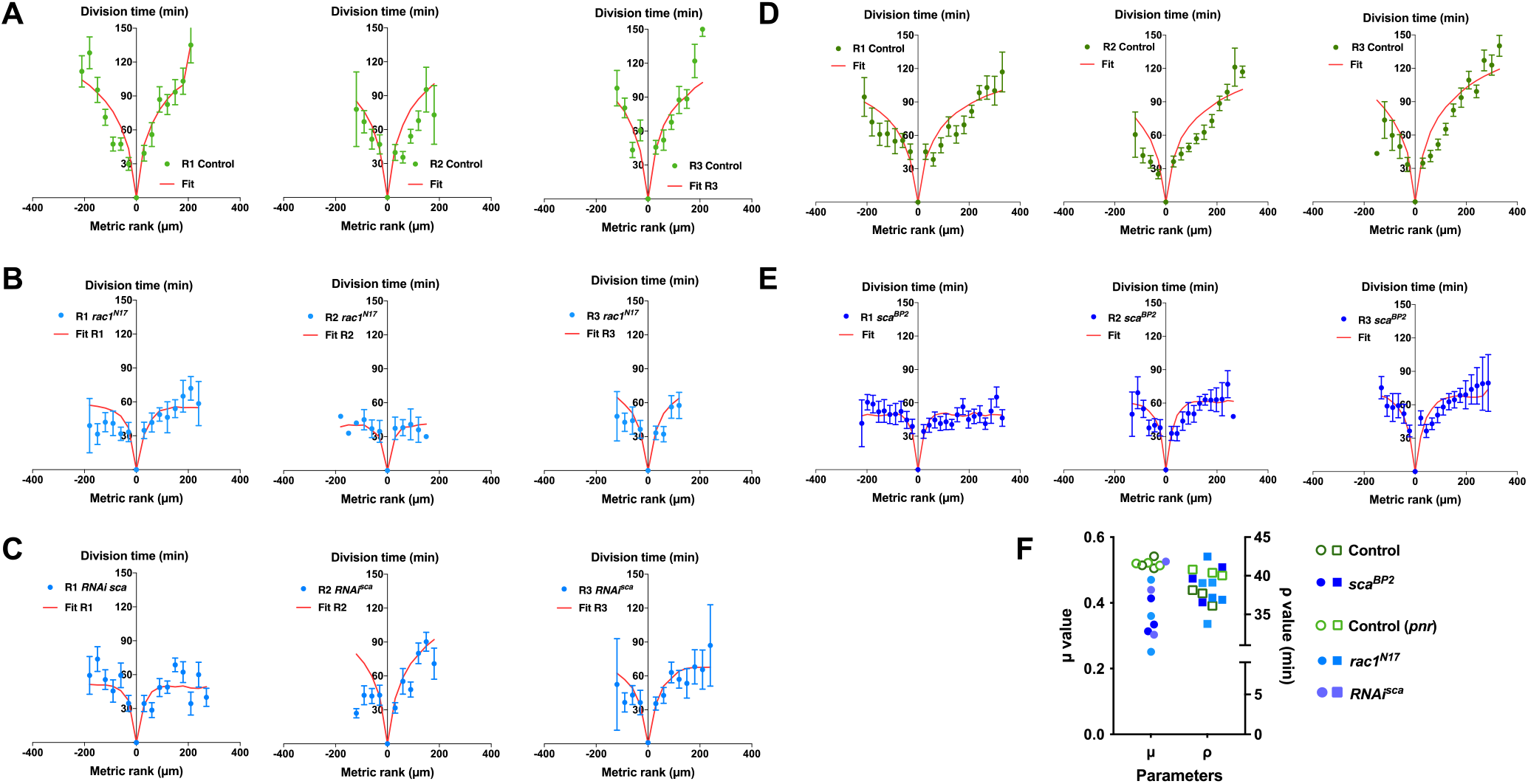
Reduced mitotic waves were better fitted with low values of the inhibitory parameter μ. (A-E) The time of SOP cell division (mean time ± SEM) in row 1 (R1), row 2 (R2) and row 3 (R3) is plotted relative to the position and time of division of the first cell to divide in each row. The different conditions in which the mitotic wave amplitude was reduced were: (B) over-expression of *rac1^N17^*, (C) conditional *RNAi-sca*, (E) *sca^BP2^* null mutant. (A) and (D) corresponding controls to B-C and E respectively. Red lines depict the best fits obtained with our model. (F) Dot plot showing the distribution μ and ρ parameters extracted from the fits displayed in A-E. Note that in all cases the best fit was obtained after the reduction of the μ parameter.

**Figure S5.**
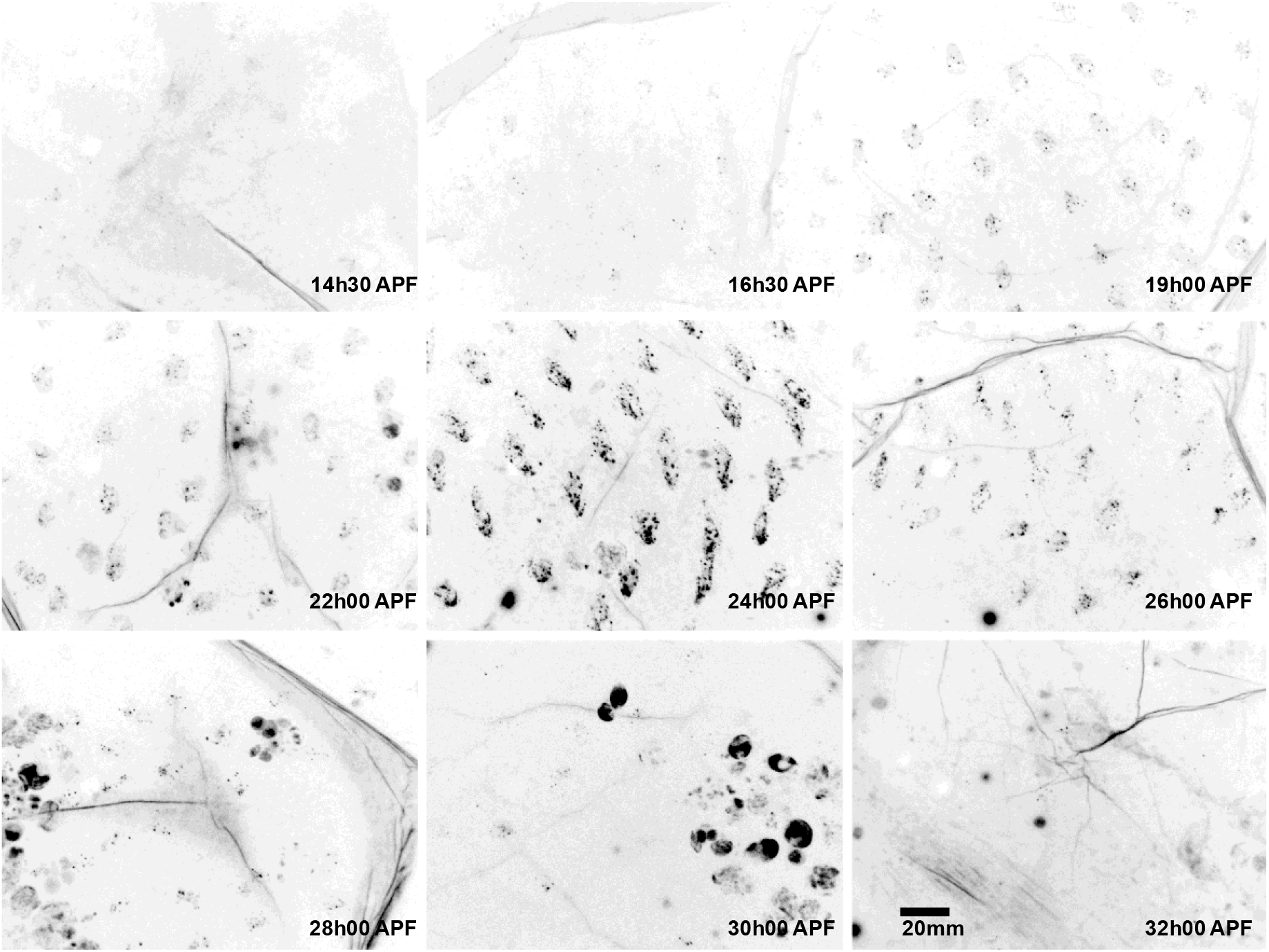
Sca is transiently expressed in the notum. Images of nota in sca::GFSTF protein-trap flies at different developmental times. Sca::GFSTF protein was transiently detected as multiple foci in sensory organ cells. Scale bar, 20 μm

**Figure S6.**
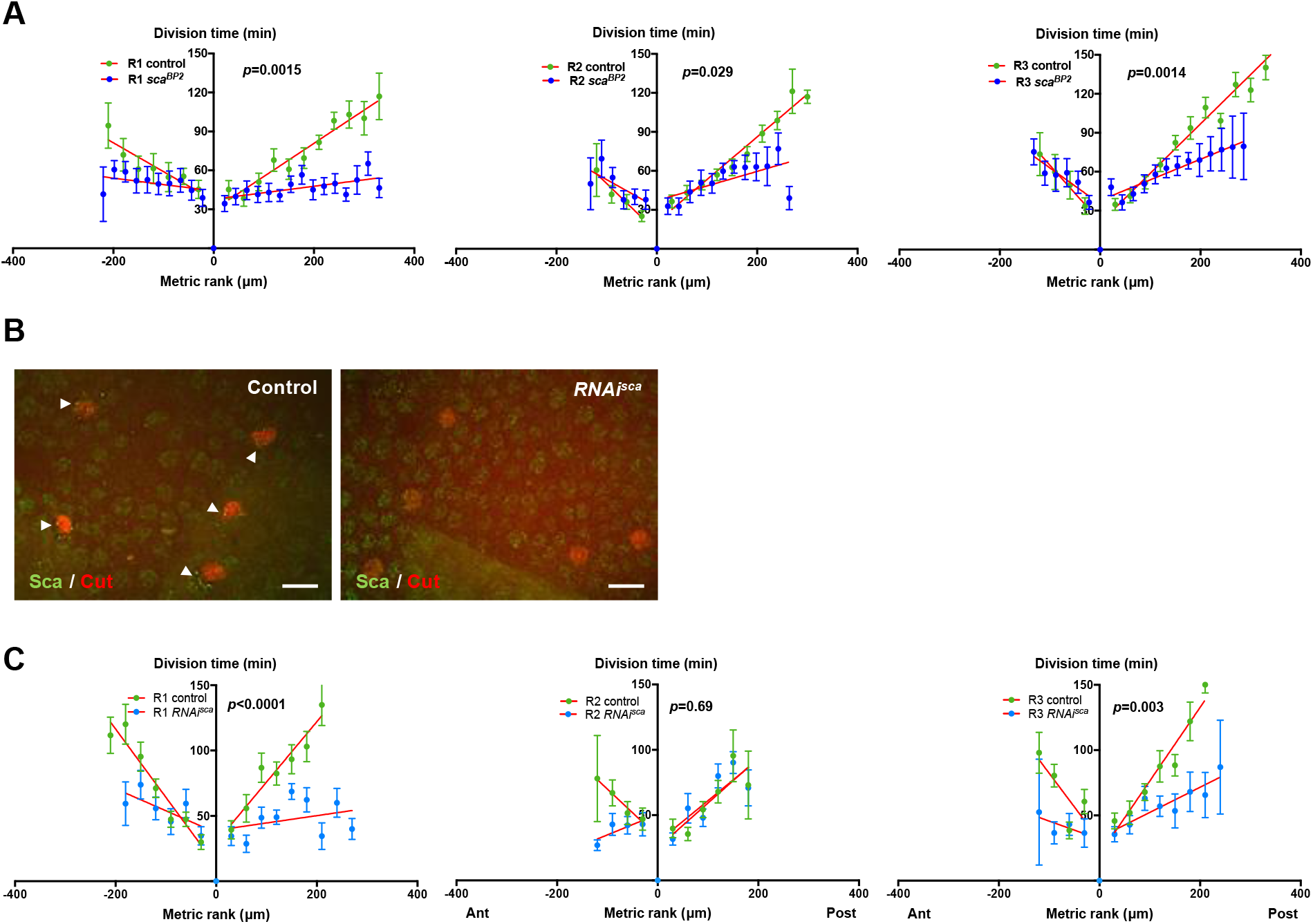
The amplitude of the mitotic wave is reduced in *sca* loss of function. (A) The relative time of SOP cell division in control and *sca^BP2^* homozygous mutant in row 1 (R1), row 2 (R2) and row 3 (R3) is plotted according to their relative position (mean time ± SEM, of 16 control nota and 9 *sca^BP2^* nota). The position and the absolute time of division of SOP_0_ in each row was taken as spatial and temporal references. The red lines correspond to standard linear regressions. (B) Efficiency of *RNAi-sca*. Immunostaining of Sca in control and after overexpression of *RNAi-sca* using the conditional genetic context *pnr>gal4 tub GAl80^ts^ RNAi-sca* in the protein trap sca::GFSTF. Pupae at 16h APF. Note that Sca spots (green, arrowheads) in SOP cells, identified by Cut staining (red), disappear in *RNAi-sca* conditions. (C) The SOP mitotic wave in control and in flies where *sca* was specifically downregulated between 12 and 19h APF, starting just before and throughout the mitotic wave, using the conditional genetic context *pnr>gal4 tub GAl80^ts^ RNAi-sca* in row 1 (R1), row 2 (R2) and row 3 (R3). Mean time ± SEM of 10 control nota and 10 *RNAi-sca* nota. The position and the absolute time of division of the first cell which divided in each row was taken as spatial and temporal reference. Red lines correspond to standard linear regressions.

**Figure S7.**
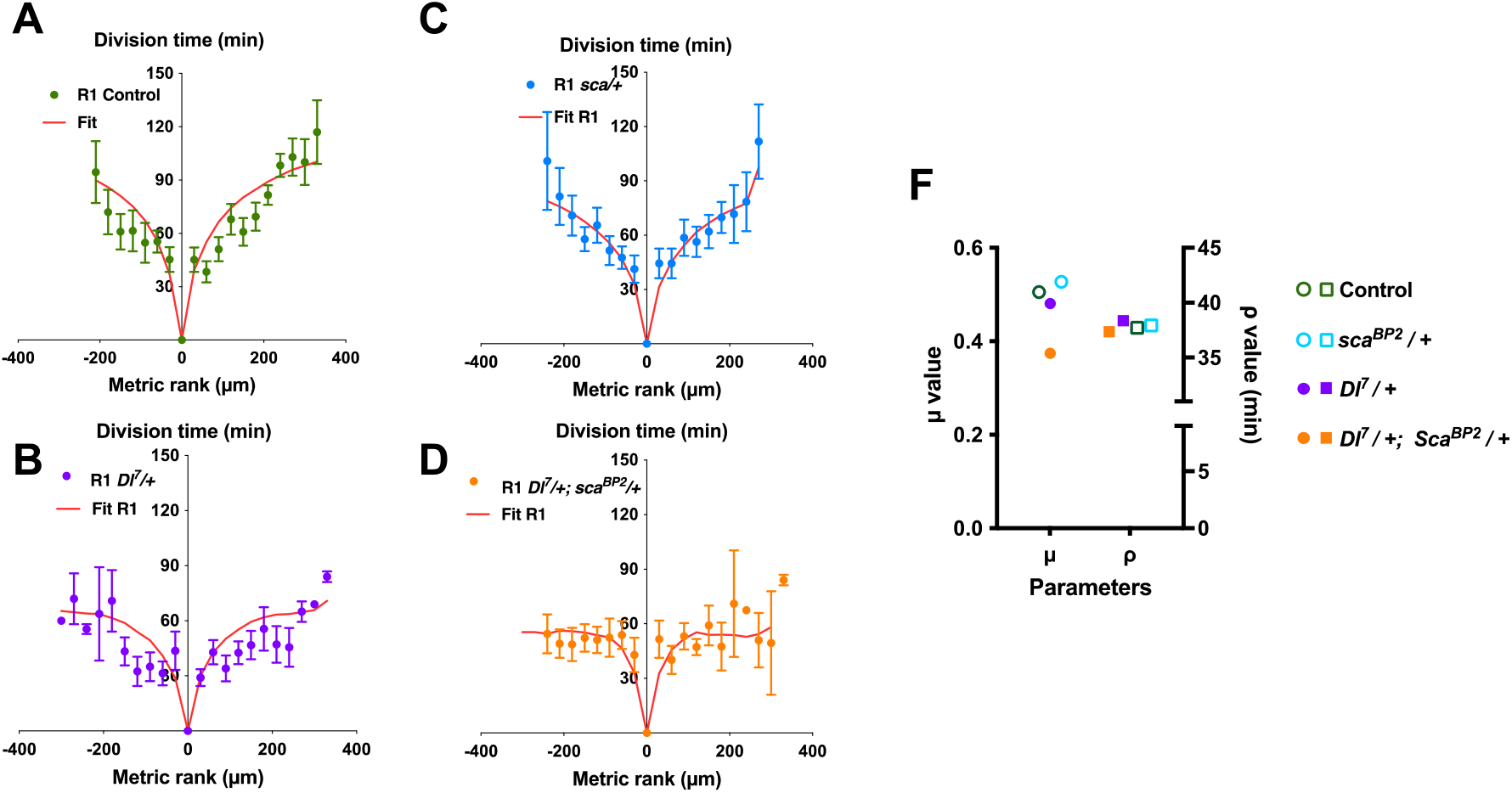
The best fit of the mitotic wave with a reduced amplitude is obtained after preferential lowering the inhibitory parameter μ. The time of SOP cell division (mean time ± SEM) in row 1 (R1), row 2 (R2) and row 3 (R3) is plotted relative to the position and time of division of the first cell to divide in each row. The different conditions were: (A) control, (B) *Dl^7^/+* heterozygous background, (C) *sca^BP2^/+* heterozygous background, (D) *Dl^7^/+, sca^BP2^/+* double heterozygous background. Red lines depict the best fits obtained with our model. (E) Dot plot showing the distribution μ and ρ parameters extracted from the fits displayed in A-D. Note that in simple heterozygous backgrounds, μ and ρ were similar to the control while in *Dl^7^/+, sca^BP2^/+* double heterozygous background, the best fit was obtained after reduction of the μ parameter only.

**Figure S8.**
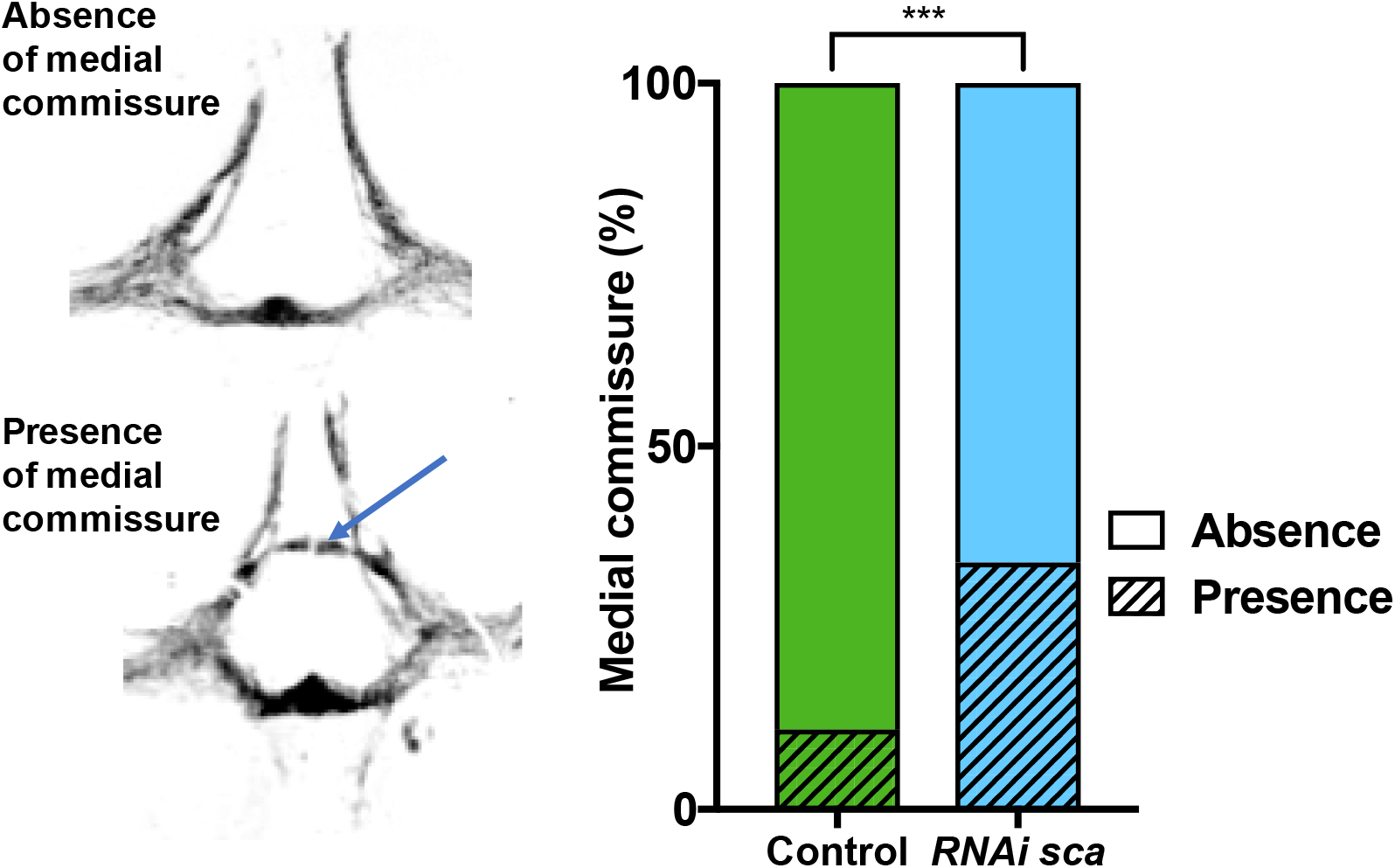
Sca downregulation modify the sensory neuropile structure. On the left, the neuropile formed by all terminal axons from sensory organs on the central notum illustrating the absence (top) and presence (bottom) of the medial commissure in the thoracic ganglia (arrow). On the right, occurrence of the medial commissure in the thoracic ganglion of control and *RNAi-sca* flies; \*\*\**p* = 0.0002, Fisher’s exact test, control flies n = 64, *RNAi-sca* flies n = 63.

## Movies

**Movie 1.** Time lapse recording of a control pupa during a 4-h period beginning at 15 h after pupal formation. The wave of SOP division is highlight in red in the row 1. Each frame was obtained by combining a z-stack (composed of 10 optical sections separated by 2 μm) acquired every 3 minutes. During the *in vivo* imaging, the pupae was maintained at 30°C. Anterior is to the left and view is dorsal. Scale bar represents 50 μm.

**Movie 2.** Time lapse recording showing the dynamic of protrusions in SOP cells in a control pupa. The membrane was visualized using a membrane tethered red fluorescent protein (RFP). Each frame was obtained by combining a z-stack (composed of 15 optical sections separated by 1 μm) acquired every 2 minutes. During the *in vivo* imaging, the pupae was maintained at 30°C. Anterior is to the left and view is dorsal. Scale bar represents 20 μm.

**Movie 3.** Time lapse recording showing the dynamics of protrusions of SOPs in pupa overexpressing the negative form of Rac 1 (*rac1^N17^*). The membrane was visualized using a membrane tethered RFP. Each frame was obtained by combining a z-stack (composed of 15 optical sections separated by 1 μm) acquired every 2 minutes. Anterior is to the left and view is dorsal. Scale bar represents 20 μm.

**Movie 4.** Time lapse showing the dynamics of Sca::GFSTF (in green) in a SOP protrusion visualized using a membrane tethered RFP (in red). Each frame was obtained by combining a z-stack (composed of 3 optical sections separated by 1 μm) acquired every 15 sec. Time in min:sec. Yellow arrowhead indicate Sca vesicle traveling along the protrusion.

**Movie 5.** Time lapse recording of the SOP mitotic wave in control pupa (top panel) and in *sca^BP2^* mutant pupa (bottom panel). For comparison, two hemi-nota are aligned. The observed difference in SOP nuclei size in the two movies is due to the lineage markers used (nuclear GFP and histone-RFP). The waves of SOP divisions are highlighted in red in the rows 1. Anterior is to the left and view is dorsal. Scale bar represents 50 μm.

**Movie 6.** Recording of cleaning reflex assay in a decapitated control adult fly. The leg movements observed were elicited after stimulation by air puffs of sensory organs in the more posterior central part of the notum. Air puffs were visualized by a red dot.

**Table S1.** Linear regression data.

| Genotype |  | Control |  |  |  |  |  |  |  |
| --- | --- | --- | --- | --- | --- | --- | --- | --- | --- |
| Linear regression | Row 1 |  | Row 2 |  | Row 3 |  | Rows 1-3 (pooled) |  | Pool (ant+post) |
|  | Anterior | Posterior | Anterior | Posterior | Anterior | Posterior | Anterior | Posterior |  |
| Best-fit values ± SE |  |  |  |  |  |  |  |  |  |
| Slope | -0.22 ± 0.046 | 0.284 ± 0.024 | -0.37 ± 0.06 | 0.3283 ± 0.0225 | -0.34 ± 0.15 | 0.3805 ± 0.0229 | -0.2631 ± 0.04339 | 0.3221 ± 0.01241 | **0.3183 ± 0.01341** |
| 1/slope | -4.5 | 3.521 | -2.7 | 3.046 | -3 | 2.628 | -3.801 | 3.105 | 3.142 |
| Speed (µm/min) | 4.5 | 3.521 | 2.7 | 3.046 | 3 | 2.628 | 3.801 | 3.105 | **3.142** |
| R square | 0.82 | 0.9342 | 0.95 | 0.9636 | 0.54 | 0.9649 | 0.8803 | 0.9854 | 0.9707 |
| Equation | Y = -0.22*X + 37 | Y = 0.284*X + 26.03 | Y = -0.37*X + 13 | Y = 0.3283*X + 20.51 | Y = -0.34*X + 26 | Y = 0.3805*X + 20.68 | Y = -0.2631*X + 27.8 | Y = 0.3221*X + 23.17 | Y = 0.3183*X + 22.9 |

| Genotype |  |  | Rac1<sup>N17</sup> |  |  |  |  |  |  |
| --- | --- | --- | --- | --- | --- | --- | --- | --- | --- |
| Linear regression | Row 1 |  | Row 2 |  | Row 3 |  | Rows 1-3 (pooled) |  | Pool (ant+post) |
|  | Anterior | Posterior | Anterior | Posterior | Anterior | Posterior | Anterior | Posterior |  |
| Best-fit values ± SE |  |  |  |  |  |  |  |  |  |
| Slope | -0.02748 ± 0.04033 | 0.1483 ± 0.03153 | -0.04935 ± 0.04737 | -0.05545 ± 0.03652 | -0.1102 ± 0.03911 | 0.3235 ± 0.127 | 0.04807 ± 0.07199 | 0.1633 ± 0.03744 | **0.1438 ± 0.03292** |
| 1/slope | -36.4 | 6.741 | -20.26 | -18.03 | -9.073 | 3.091 | 20.8 | 6.123 | 6.952 |
| Speed (µm/min) | 36.4 | 6.741 | 20.26 | 18.03 | 9.073 | 3.091 | 27.25 | 6.123 | **6.952** |
| R square | 0.104 | 0.7867 | 0.2134 | 0.4346 | 0.7988 | 0.1603 | 0.446 | 0.1252 | 0.7367 |
| Equation | Y = -0.02748*X + 33.54 | Y = 0.1483*X + 32.61 | Y = -0.04935*X + 34.76 | Y = -0.05545*X + 41.41 | Y = -0.1102*X + 34.55 | Y = 0.3235*X + 20.39 | Y = 0.04807*X + 34.82 | Y = 0.1633*X + 29.77 | Y = 0.1438*X + 30.03 |

| Genotype |  | Sca<sup>RNAi</sup> |  |  |  |  |  |  |  |
| --- | --- | --- | --- | --- | --- | --- | --- | --- | --- |
| Linear regression | Row 1 |  | Row 2 |  | Row 3 |  | Rows 1-3 (pooled) |  | Pool (ant+post) |
|  | Anterior | Posterior | Anterior | Posterior | Anterior | Posterior | Anterior | Posterior |  |
| Best-fit values ± SE |  |  |  |  |  |  |  |  |  |
| Slope | -0.1686 ± 0.08467 | 0.0561 ± 0.06099 | 0.1576 ± 0.09048 | 0.3172 ± 0.1109 | -0.1383 ± 0.09706 | 0.1929 ± 0.04038 |  |  |  |
| 1/slope | -5.932 | 17.83 | 6.346 | 3.152 | -7.233 | 5.185 |  |  |  |
| Speed (µm/min) | 5.932 | 17.83 | 6.346 | 3.152 | 7.233 | 5.185 |  |  |  |
| R square | 0.4978 | 0.1078 | 0.6027 | 0.6716 | 0.5036 | 0.7917 | na |  |  |
| Equation | Y = -0.1686*X + 37.1 | Y = 0.0561*X + 38.95 | Y = 0.1576*X + 50.6 | Y = 0.3172*X + 29.36 | Y = -0.1383*X + 31.81 | Y = 0.1929*X + 32.99 |  |  |  |

| Genotype |  | *sca*<sup>BP2</sup>/*sca*<sup>BP2</sup> |  |  |  |  |  |  |  |
| --- | --- | --- | --- | --- | --- | --- | --- | --- | --- |
| Linear regression | Row 1 |  | Row 2 |  | Row 3 |  | Rows 1-3 (pooled) |  | Pool (ant+post) |
|  | Anterior | Posterior | Anterior | Posterior | Anterior | Posterior | Anterior | Posterior |  |
| Best-fit values ± SE |  |  |  |  |  |  |  |  |  |
| Slope | -0.04933 ± 0.03233 | 0.04986 ± 0.01626 | -0.2155 ± 0.1066 | 0.1123 ± 0.04281 | -0.2776 ± 0.06459 | 0.1587 ± 0.01431 |  |  |  |
| 1/slope | -20.27 | 20.06 | -4.641 | 8.902 | -3.603 | 6.303 |  |  |  |
| Speed (µm/min) | 20.27 | 20.06 | 4.641 | 8.902 | 3.603 | 6.303 |  |  |  |
| R square | 0.2254 | 0.4197 | 0.5052 | 0.4078 | 0.8219 | 0.9179 |  |  | na |
| Equation | Y = -0.04933*X + 44.16 | Y = 0.04986*X + 37.78 | Y = -0.2155*X + 31.71 | Y = 0.1123*X + 37.13 | Y = -0.2776*X + 35.15 | Y = 0.1587*X + 37.87 |  |  |  |

| Genotype |  | *DI*<sup>7</sup>/+ |  |
| --- | --- | --- | --- |
| Linear regression | Row 1 |  | Pool (ant+post) |
|  | Anterior | Posterior |  |
| Best-fit values ± SE |  |  |  |
| Slope | -0.1325 ± 0.03879 | 0.1448 ± 0.02417 | 0.1472 ± 0.02304 |
| 1/slope | -7.548 | 6.906 | 6.791 |
| Speed (µm/min) | 7.548 | 6.906 | 6.791 |
| R square | 0.5932 | 0.7995 | 0.8195 |
| Equation | Y = -0.1325*X + 28.93 | Y = 0.1448*X + 25 | *Y = 0.1472*X + 25.93* |

| Genotype |  | *sca*<sup>BP2/+</sup> |  |
| --- | --- | --- | --- |
| Linear regression | Row 1 |  | Pool (ant+post) |
|  | Anterior | Posterior |  |
| Best-fit values ± SE |  |  |  |
| Slope | -0.2487 ± 0.05619 | 0.2285 ± 0.03891 | 0.2547 ± 0.02863 |
| 1/slope | -4.022 | 4.377 | 3.925 |
| Speed (µm/min) | 4.022 | 4.377 | 3.925 |
| R square | 0.1488 | 0.8312 | 0.9188 |
| Equation | Y = -0.2487*X + 31.15 | Y = 0.2285*X + 32.13 | Y = 0.2547*X + 30.04 |

| Genotype |  | *DI*<sup>7</sup>/*sca*<sup>BP2</sup> |  |
| --- | --- | --- | --- |
| Linear regression | Row 1 |  | Pool (ant+post) |
|  | Anterior | Posterior |  |
| Best-fit values ± SE |  |  |  |
| Slope | -0.01871 ± 0.01919 | 0.03806 ± 0.03443 | 0.02307 ± 0.01829 |
| 1/slope | -53.45 | 26.27 | 43.35 |
| Speed (µm/min) | 53.45 | 26.27 | 43.35 |
| R square | 0.9508 | 0.1325 | 0.1659 |
| Equation | Y = -0.01871*X + 48.08 | Y = 0.03806*X + 47.49 | Y = 0.02307*X + 48.35 |

**Table S2.** Values of ρ and μ parameters obtained after better fitting for all experimental conditions.

| Genotype | Row | μ | ρ |
| --- | --- | --- | --- |
| Control *Sca*<sup>BP2</sup> | 1 | 0,5051027 | 37,70738036 |
|  | 2 | 0,51370006 | 36,11782771 |
|  | 3 | 0,5418 | 38,136 |
| *Sca*<sup>BP2</sup> | 1 | 0,31375 | 41,087 |
|  | 2 | 0,4138459 | 36,56211578 |
|  | 3 | 0,3340962 | 39,6176644 |
| Control *rac1*<sup>N17</sup> | 1 | 0,52179866 | 40,38357955 |
|  | 2 | 0,52572031 | 40,02894363 |
|  | 3 | 0,51446082 | 40,80249035 |
| *Rac1*<sup>N17</sup> | 1 | 0,36045363 | 39,12344446 |
|  | 2 | 0,2510631 | 39,04736903 |
|  | 3 | 0,47042889 | 33,78043964 |
| Control *sca*<sup>RNAi</sup> | 1 | 0,52179866 | 40,38357955 |
|  | 2 | 0,52572031 | 40,02894363 |
|  | 3 | 0,51446082 | 40,80249035 |
| *Sca*<sup>RNAi</sup> | 1 | 0,30340662 | 42,48175604 |
|  | 2 | 0,51970364 | 37,15218064 |
|  | 3 | 0,43959156 | 36,89061269 |
| *sca*<sup>BP2</sup>/+ | 1 | 0,52731456 | 37,9225975 |
| *DI*/+ | 1 | 0,48094919 | 38,34110558 |
| *sca*<sup>BP2</sup>/+; *DI* / + | 1 | 0,37443512 | 37,35840387 |

## Key resource table

| Reagent or resource | source | identifier | Additional information |
| --- | --- | --- | --- |
| **Antibodies** |  |  |  |
| mouse anti-Futsch (clone 22C10) | DSHB | AB_528403 | 1/200 |
| rabbit anti-HA (clone 3F10) | Roche | AB_231 4622 | 1/250 |
| mouse anti-Flag (M2) | Sigma | AB_259529 | 1/250 |
| anti-V5 (E10/V4RR), DyLight650 | ThermoFisher | AB_2537642 | 1/100 |
| anti-cut | DSHB | AB_528186 | 1/500 |
| **Fly strains** |  |  |  |
| *w<sup>1118</sup>* | Bloomington Stock Center | BDSC_3605 |  |
| *neurD>GFP* | Schweisguth F. |  |  |
| *sca<sup>BP2</sup>* | Bloomington Stock Center | BDSC_7320 |  |
| *UAS sca-RNAi* | Vienna Drosophila Resource | VDRC#44527 |  |
| *sca::GFSTF* | Bloomington Stock Center | BDSC_64443 |  |
| *neur>ph (PLCd)::RFP* | Schweisguth F. |  |  |
| *UAS-MCFO-1* | Bloomington Stock Center | BDSC_64085 |  |
| *UAS-rac1<sup>N17</sup>* |  | BDSC_6292 |  |
| *DI<sup>7</sup>* |  | BDSC_485 |  |
| *UAS-ph::GFP* | Bellaiche Y. |  |  |
| *neur<sup>p72</sup>>Gal4* | Bellaiche Y. |  |  |

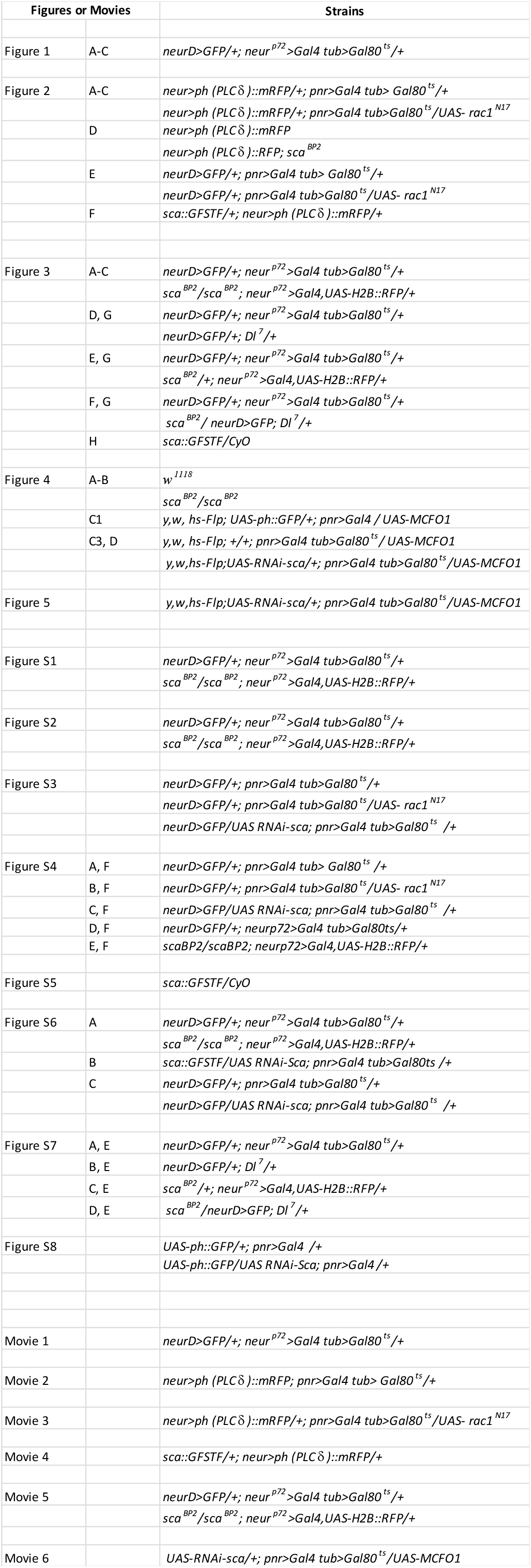

## REFERENCES

1. Holguera, I., and Desplan, C. (2018). Neuronal specification in space and time. Science 362, 176–180.

2. Hirata, T., and Iwai, L. (2019). Timing matters: A strategy for neurons to make diverse connections. Neurosci. Res. 138, 79–83.

3. Gatto, G., Smith, K.M., Ross, S.E., and Goulding, M. (2019). Neuronal diversity in the somatosensory system: bridging the gap between cell type and function. Curr. Opin. Neurobiol. 56, 167–174.

4. Sato, M., Kojima, T., Michiue, T., and Saigo, K. (1999). Bar homeobox genes are latitudinal prepattern genes in the developing Drosophila notum whose expression is regulated by the concerted functions of decapentaplegic and wingless. Development 126, 1457–1466.

5. Usui, K., and Kimura, K. (1993). Sequential emergence of the evenly spaced microchaetes on the notum of Drosophila. Rouxs Arch. Dev. Biol. 203, 151–158.

6. Corson, F., Couturier, L., Rouault, H., Mazouni, K., and Schweisguth, F. (2017). Self-organized Notch dynamics generate stereotyped sensory organ patterns in Drosophila. Science 356, eaai7407.

7. Gho, M., Bellaïche, Y., and Schweisguth, F. (1999). Revisiting the Drosophila microchaete lineage. Development 126, 3573–3584.

8. Ghysen, A. (1980). The projection of sensory neurons in the central nervous system of Drosophila: Choice of the appropriate pathway. Dev. Biol. 78, 521–541.

9. Usui-Ishihara, A., and Simpson, P. (2005). Differences in sensory projections between macro- and microchaetes in Drosophilid flies. Dev. Biol. 277, 170–183.

10. Corfas, G., and Dudai, Y. (1989). Habituation and dishabituation of a cleaning reflex in normal and mutant Drosophila. J. Neurosci. 9, 56–62.

11. Mok, L.-P., Qin, T., Bardot, B., LeComte, M., Homayouni, A., Ahimou, F., and Wesley, C. (2005). Delta activity independent of its activity as a ligand of Notch. BMC Dev. Biol. 5, 6.

12. Renaud, O., and Simpson, P. (2001). scabrous Modifies Epithelial Cell Adhesion and Extends the Range of Lateral Signalling during Development of the Spaced Bristle Pattern in Drosophila. Dev. Biol. 240, 361–376.

13. Lee, E.-C., Yu, S.-Y., and Baker, N.E. (2000). The Scabrous protein can act as an extracellular antagonist of Notch signaling in the Drosophila wing. Curr. Biol. 10, 931–S2.

14. Li, Y. (2003). Scabrous and Gp150 are endosomal proteins that regulate Notch activity. Development 130, 2819–2827.

15. Muñoz-Soriano, V., Santos, D., Durupt, F.C., Casani, S., and Paricio, N. (2016). Scabrous overexpression in the eye affects R3/R4 cell fate specification and inhibits notch signaling: SCA Overexpression Affects R3/R4 Cell Specification. Dev. Dyn. 245, 166–174.

16. Powell, P.A., Wesley, C., Spencer, S., and Cagan, R.L. (2001). Scabrous complexes with Notch to mediate boundary formation. Nature 409, 626–630.

17. Cohen, M., Georgiou, M., Stevenson, N.L., Miodownik, M., and Baum, B. (2010). Dynamic Filopodia Transmit Intermittent Delta-Notch Signaling to Drive Pattern Refinement during Lateral Inhibition. Dev. Cell 19, 78–89.

18. Hunter, G.L., Hadjivasiliou, Z., Bonin, H., He, L., Perrimon, N., Charras, G., and Baum, B. (2016). Coordinated control of Notch/Delta signalling and cell cycle progression drives lateral inhibition-mediated tissue patterning. Development 143, 2305–2310.

19. Buszczak, M., Inaba, M., and Yamashita, Y.M. (2016). Signaling by Cellular Protrusions: Keeping the Conversation Private. Trends Cell Biol. 26, 526–534.

20. Kornberg, T.B., and Roy, S. (2014). Cytonemes as specialized signaling filopodia. Development 141, 729–736.

21. Baker, N., Mlodzik, M., and Rubin, G. (1990). Spacing diffentiation in the developing Drosophila eye: A fibrinogen-related lateral inhibitor encoded by scabrous. Science 250, 1370–1377.

22. Gavish, A., Shwartz, A., Weizman, A., Schejter, E., Shilo, B.-Z., and Barkai, N. (2016). Periodic patterning of the Drosophila eye is stabilized by the diffusible activator Scabrous. Nat. Commun. 7.

23. Chou, Y.-H., and Chien, C.-T. (2002). Scabrous Controls Ommatidial Rotation in the Drosophila Compound Eye. Dev. Cell 3, 839–850.

24. Petruccelli, E., Feyder, M., Ledru, N., Jaques, Y., Anderson, E., and Kaun, K.R. (2018). Alcohol Activates Scabrous-Notch to Influence Associated Memories. Neuron 100, 1209–1223.e4.

25. Nern, A., Pfeiffer, B.D., and Rubin, G.M. (2015). Optimized tools for multicolor stochastic labeling reveal diverse stereotyped cell arrangements in the fly visual system. Proc. Natl. Acad. Sci. 112, E2967–E2976.

26. Vandervorst, P., and Ghysen, A. (1980). Genetic control of sensory connections in Drosophila. Nature 286, 65–67.

27. Otsuki, L., and Brand, A.H. (2018). Cell cycle heterogeneity directs the timing of neural stem cell activation from quiescence. Science 360, 99–102.

28. Fernandes, V.M., Chen, Z., Rossi, A.M., Zipfel, J., and Desplan, C. (2017). Glia relay differentiation cues to coordinate neuronal development in Drosophila. Science 357, 886–891.

29. Buffin, E., and Gho, M. (2010). Laser microdissection of sensory organ precursor cells of Drosophila microchaetes. PloS One 5, e9285.

30. Muskavitch, M.A. (1994). Delta-notch signaling and Drosophila cell fate choice. Dev. Biol. 166, 415–430.

31. Ayeni, J.O., Audibert, A., Fichelson, P., Srayko, M., Gho, M., and Campbell, S.D. (2016). G2 phase arrest prevents bristle progenitor self-renewal and synchronizes cell division with cell fate differentiation. Development 143, 1160–1169.

32. Fichelson, P., and Gho, M. (2004). Mother–daughter precursor cell fate transformation after Cdc2 down-regulation in the Drosophila bristle lineage. Dev. Biol. 276, 367–377.

33. Deng, W.-M., Althauser, C., and Ruohola-Baker, H. (2001). Notch-Delta signaling induces a transition from mitotic cell cycle to endocycle in Drosophila follicle cells. Development 128, 4737–4746.

34. Krejčí, A., Bernard, F., Housden, B.E., Collins, S., and Bray, S.J. (2009). Direct Response to Notch Activation: Signaling Crosstalk and Incoherent Logic. Sci. Signal. 2, ra1–ra1.

35. Aradhya, R., Zmojdzian, M., Da Ponte, J.P., and Jagla, K. (2015). Muscle niche-driven Insulin-Notch-Myc cascade reactivates dormant Adult Muscle Precursors in Drosophila. eLife 4.

36. McCormick, L.E., and Gupton, S.L. (2020). Mechanistic advances in axon pathfinding. Curr. Opin. Cell Biol. 63, 11–19.

37. Petrovic, M., and Hummel, T. (2008). Temporal identity in axonal target layer recognition. Nature 456, 800–803.

38. Bellaïche, Y., Gho, M., Kaltschmidt, J.A., Brand, A.H., and Schweisguth, F. (2001). Frizzled regulates localization of cell-fate determinants and mitotic spindle rotation during asymmetric cell division. Nat. Cell Biol. 3, 50–57.

39. Sato, M., and Saigo, K. (2000). Involvement of pannier and u-shaped in regulation of Decapentaplegic-dependent wingless expression in developing Drosophila notum. Mech. Dev. 93, 127–138.

40. Couturier, L., Vodovar, N., and Schweisguth, F. (2012). Endocytosis by Numb breaks Notch symmetry at cytokinesis. Nat. Cell Biol. 14, 131–139.

41. Schindelin, J., Arganda-Carreras, I., Frise, E., Kaynig, V., Longair, M., Pietzsch, T., Preibisch, S., Rueden, C., Saalfeld, S., Schmid, B., et al. (2012). Fiji: an open-source platform for biological-image analysis. Nat. Methods 9, 676–682.

