## Supplementary material for "A neural progenitor mitotic wave is required for asynchronous axon outgrowth and morphology": Model Description

LACOSTE ET AL.

### 1. Phenomenological mode description

In this section, we describe a very simple model to explain the the shape of the wave. Let's have $k \geq 0$ the cells' indices. Cell's have a waiting time to divide $t_k$ this waiting time is the time when some cell's value $x_k$ reach a constant threshold $\theta$ assumed equal for all cells. Therefore, $t_k$ is the time when $x_k(t_k) = \theta$.

In order to reach $\theta$ we assume that cells have two speeds : one $v$ is when the cell is not inhibited. Starting from 0 a non inhibited cell take $\theta/v$ time to divide. When cells are inhibited the speed is reduced to $\lambda v$ with $0 \leq \lambda \leq 1$. A value $\lambda$ close to 0 is a strong inhibition a value close to 1 is a light inhibition.

Assuming at first that all cells except cell $k = 0$ are inhibited and that cells inhibit their next neighbor $k + 1$ and are inhibited by their previous neighbor $k - 1$. We can compute the time to division for cell 0 since it is not inhibited: $\theta/v$.

For cell $k$ we have the following relationship:

$$x_k(t_k) = \theta = \lambda v t_{k-1} + v\left(t_k - t_{k-1}\right)$$

Since the cell $k$ has been inhibited until the cell $k-1$ has divided, the time for division is the time from the start with neighbor inhibition $\lambda v t_{k-1}$ and then since the cell $k-1$ does not inhibit anymore after $t_{k-1}$ the remaining time to the threshold is $v\left(t_k - t_{k-1}\right)$ at full speed.

We can rearrange:

$$\begin{aligned}
\theta &= \lambda v t_{k-1} + v\left(t_k - t_{k-1}\right) \\
v t_k &= \theta - \lambda v t_{k-1} + v t_{k-1} \\
t_k &= a t_{k-1} + b
\end{aligned}$$

with $a = 1 - \lambda$ and $b = \theta/v = t_0$. This sequence can be solved exactly:

$$t_k = (1-\lambda)^k \left(\frac{\theta}{v} - \frac{\theta}{\lambda v}\right) + \frac{\theta}{\lambda v}$$

With $\lambda$ close to 1 - no inhibition - we obtain $t_k = b = \theta/v$ every cells divides at the same time.

For low values of $\lambda$ - high inhibition - the curve display a wave of consecutive divisions. At the limit of total inhibition $\lambda = 0$ the division time is a line $t_k = bk = k\theta/v$.

If we take $v = 1$ and $\lambda = 0.5$ we have:

$$\begin{aligned} t_0 &= 1 \\ t_1 &= 1.5 \\ t_2 &= 1.75 \\ t_3 &= 1.875 \end{aligned}$$

In this example, we have a propagating wave of division that accelerates through time flattening the curve. In case of no inhibition the curve is almost flat passes the first division.

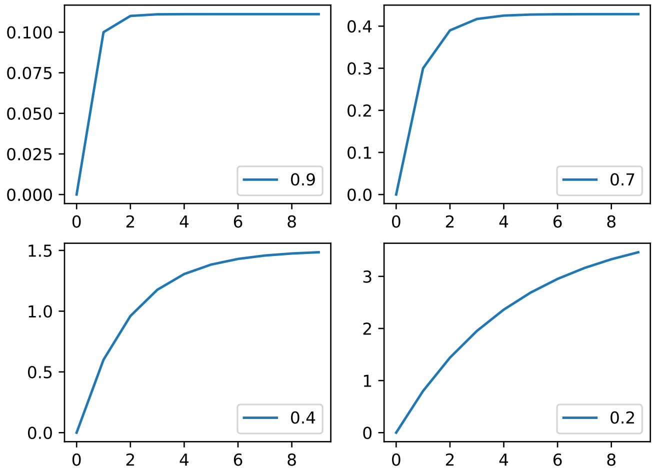


FIGURE 1. Example of the phenomenological model with $\theta = 1$ and $v = 1$. $t_k - t_0$ for $k \in \{0, \ldots, 9\}$. For $\lambda \in \{.9, 0.7, 0.4, 0.2\}$ for low to strong inhibition. Please note that the y scales are different.

### 2. Model description

In order to compare with the experimental data, we developed another mathematical model. In this model we make similar assumptions: we first assume that (1) neighboring progenitor cells are in a row and inhibit each other to enter into mitosis through cell contacts and (2) this inhibition is switched off when cells divide. We also assume that (3) cell division occurs whenever a certain cellular compound concentration (called $A$) reaches a certain threshold ($\theta$). Finally, (4) this compound is produced at a constant rate but this production is inhibited (decreased) by direct neighbors cells thus preventing the current cell from dividing.

Using dynamical systems, this can be modeled as (for a given cell indexed by $k$):

$$\rho \frac{dA_k}{dt} = \frac{r}{1+I_k} - A_k \quad (1)$$

$$\delta \frac{dI_k}{dt} = \mu \, \mathrm{H}\,(A_{k+1}-\theta) + \mu \, \mathrm{H}\,(A_{k-1}-\theta) - I_k \quad (2)$$

with

$$\mathrm{H}(x) = \begin{cases} 1 & x < 0 \\ 0 & x \geq 0 \end{cases}$$

As mentioned $A$ describes the compound concentration with constant production rate ($r$) inhibited by $I$ another compound that decreases $A$'s total production rate. All degradations are at a constant rate ($\rho$ and $\delta$) as well. Inhibition is computed cell-wise and depends on neighbors' status. The production of inhibition is driven by $\mu$. Higher values of $\mu$ indicate greater inhibition from neighboring cells. Neighboring cells whose $A$ is below the threshold (i.e that are not dividing) maintain the inhibition. Whenever the cell divides, inhibition vanishes. Please note that the extent of the inhibition is for direct neighbors only hence the $+1$ and $-1$ in the index.

We can simulate variability in this system of equations by adding a random noise in the starting values of $A$.

This figure shows the main features of the model: to obtain similar division rates with successive cell divisions (a linear increasing division time function) a strong inhibition is needed. On the other hand, almost simultaneous cells division (without any wave) is obtained by decreasing the amount of inhibition.

### 3. Parameters estimation

To estimate parameters, we perform $L_2$ norm minimization :

$$\chi^2(r,\rho,\mu,\delta) = \Sigma_{k=0}^{N} \left(T_k - t_k(r,\rho,\mu,\delta)\right)^2 \quad (3)$$

$T_k$ are the division time for position $k \in \{\ldots,-3,-2,-1,0,1,2,3,\ldots\}$ from the data and $t_k(r,\rho,\mu,\delta)$ is the division time from the model with parameters $r,\rho,\mu,\delta$. We performed Nelder-Mead parameters optimisations in order to reach minimal $\chi^2$.

In this model, all parameters cannot be identified exactly at once since the only data fitted is the division time: having four parameters has produced redundant information to fit the division time from the data. To avoid problems of identification, we assumed that production rate $r$ is unlikely to change. We assumed also that degradation for the inhibitory compound was higher therefore their half-life constant much lower ($\delta$ is small compared to $\rho$) and also set it to a constant. These assumptions allows to reduce the problem to two free parameters - $\mu$ and $\rho$ - to perform the fitting. In this case, decreasing the number of free parameters had no significant impact in $L_2$ minimisation as fits were comparable while maintaining stable results.

In order to obtain noisy representations, the algorithm performs 10 simulation of the differential equation with random starting values for $A$ (uniform distribution between 0 and

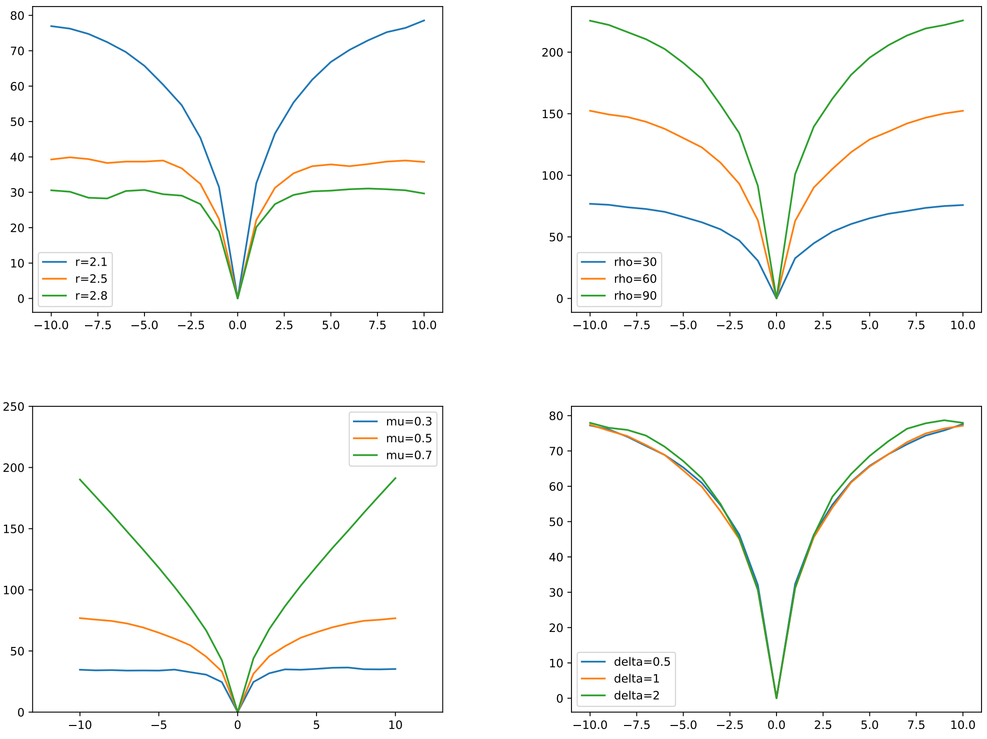


FIGURE 2. TOP LEFT: Simulation different production rate $r \in \{2.1, 2.5, 2.8\}$ (other parameters are $\rho = 30$, $\mu = 0.5$, $\delta = 1$). TOP RIGHT: Simulation with various temporal constant $\rho \in \{30, 60, 90\}$ (other parameters are $r = 2.1$, $\mu = 0.5$, $\delta = 1$). This adjusts the temporal scale but not the shape of the wave BOTTOM LEFT simulation with various inhibition strength with $\mu \in \{0.5, 0.6, 0.7\}$ (other parameters are $r = 2.1$, $\rho = 30$, $\delta = 1$). Strong inhibition here (0.7) imposes a more linear division time shape whereas low inhibition allows for quicker division with neighboring cells dividing almost simultaneously. BOTTOM RIGHT: simulation with various inhibition time constant with $\delta \in \{10, 0.5, 1\}$ (other parameters are $r = 2.1$, $\rho = 30$, $\mu = 0.5$).

$\theta/2$) except for the cell at position 0 which is set above threshold. The times of division are computed for each cell and the average is used as values $t_k$.
